# Wikipedia network analysis of cancer interactions and world influence

**DOI:** 10.1101/527879

**Authors:** Guillaume Rollin, José Lages, Dima L. Shepelyansky

## Abstract

We apply the Google matrix algorithms for analysis of interactions and influence of 37 cancer types, 203 cancer drugs and 195 world countries using the network of 5 416 537 English Wikipedia articles with all their directed hyperlinks. The PageRank algorithm provides a ranking of cancers which has 60% and 70% overlaps with the top 10 deadliest cancers extracted from World Health Organization GLOBOCAN 2018 and Global Burden of Diseases Study 2017, respectively. The recently developed reduced Google matrix algorithm gives networks of interactions between cancers, drugs and countries taking into account all direct and indirect links between these selected 435 entities. These reduced networks allow to obtain sensitivity of countries to specific cancers and drugs. The strongest links between cancers and drugs are in good agreement with the approved medical prescriptions of specific drugs to specific cancers. We argue that this analysis of knowledge accumulated in Wikipedia provides useful complementary global information about interdependencies between cancers, drugs and world countries.

## Introduction

“Nearly every family in the world is touched by cancer, which is now responsible for almost one in six deaths globally” [1]. The number of new cancer cases in the world is steadily growing reaching 18.1 million projected for 2018 [2] with predicted new cases of 29.4 million for 2035 [3]. The detailed statistical analysis of new cases and mortality projected for 2018 is reported in [4]. Such statistical analysis is of primary importance for estimating the influence of cancer diseases on the world population. However, it requires significant efforts of research groups and medical teams all over the world such as consortia involved in the Global Burden of Diseases Study (GBD) [5] and the WHO GLOBOCAN reports [2].

Here we propose to probe the network of Wikipedia articles in order to infer specific interactions between cancer types and to measure world influence of cancers. Wikipedia can be seen as a global database of accumulated human knowledge with an immense variety of topics. Moreover the way Wikipedia articles are citing each other encodes scientific, social, historical, and many other aspects. In principle one should be able to extract from Wikipedia direct or indirect relations between cancers, drug cancers and countries. We focus our study on cancer which is one of the major cause of human mortality and which consequently have important social and political impacts all around the world. We aim to measure these impacts through the prism of Wikipedia.

Thus we develop a complementary approach to the existing statistical approaches [2, 4, 5], the Wikipedia network analysis based on the Google matrix and PageRank algorithm invented by Brin and Page in 1998 for World Wide Web search engine information retrieval [6, 7]. Applications of this approach to various directed networks are described at [8]. Here we use the network of English Wikipedia articles collected in May 2017 with *N* = 5 416 537 articles and connected by *N*_*l*_ = 122 232 032 directed links, i.e. quotations from one article to another.

At present Wikipedia represents a public, open, collectively created encyclopaedia with a huge amount of information exceeding those of Encyclopedia Britannica [9] in volume and accuracy of articles devoted to scientific topics [10]. As an example, articles on biomolecules are actively maintained by Wikipedians [11, 12]. The academic analysis of information collected in Wikipedia is growing, getting more tools and applications as reviewed in [13, 14]. The scientific analysis shows that the quality of Wikipedia articles is growing [15].

A new element of our analysis is the reduced Google matrix (REGOMAX) method developed recently [16, 17]. This method selects a modest size subset of *N*_*r*_ nodes of interest from a huge global directed network with *N* ≫ *N*_*r*_ nodes and generates the reduced Google matrix *G*_R_ taking into account all direct pathways and indirect pathways (i.e. those going through the global network) between the *N*_*r*_ nodes. This approach conserves the PageRank probabilities of nodes from the global Google matrix *G* (up to a normalization factor). This method uses the ideas coming from the scattering theory of complex nuclei, mesoscopic physics and quantum chaos.

The efficiency of this approach has been tested with Wikipedia networks of politicians [17], painters [18], world universities [19], with biological networks from SIGNOR data base [20], with world trade networks [21, 22], and with financial networks [23]. The method is general as it can be applied to any subset of nodes embedded in a huge directed network. The main outcome is a synthetic effective view of the subnetwork encoded by weighted links in the corresponding reduced Google matrix. The strength of the specific application of REGOMAX method to Wikipedia networks is the encyclopedic nature of Wikipedia. Myriads of subjects are treated in Wikipedia which allow through the network of articles to connect, at least indirectly, many very different topics such as, e.g., for the present study, countries and cancer types. Moreover in the framework of the REGOMAX method every articles in Wikipedia, even articles having apparently nothing to do with the subjects of interest possibly contribute to the effective link obtained between two chosen nodes (i.e., articles). Although the quality of Wikipedia articles is constantly growing [10–12, 15], the information extraction may be sensitive to noise coming from inadequate or not so relevant links introduced in certain Wikipedia articles. These noisy links, which depends on when the Wikipedia network has been extracted, usually are cleaned out by the Wikipedians collaborative effort. The lifetime before removal of these noisy links depend also on the subject. As there is no simple way to quantify this source of noise (the degree of relevance of a link between two articles), we assume that in average it causes no harm to the present study. The results are presented in the devoted section keeping in mind this limitation.

In this work the reduced network is composed of *N*_*cr*_ = 37 types of cancers listed at Wikipedia [24] and *N*_*d*_ = 203 drugs for cancer extracted from data base [25]. All these *N*_*cr*_ + *N*_*d*_ = 240 items had an active Wikipedia article in May 2017. All these cancers and drugs are listed in alphabetic order in Tabs. 1 and 2. In addition we add to the selected set of articles *N*_*cn*_ = 195 world countries that allows us to analyze the global influence of cancer types (the ranking and REGOMAX analysis of countries are reported in [26, 27]). The PageRank list of the 195 selected countries is available at [28]. Thus in total the reduced Google matrix selected number of nodes is *N*_*r*_ = *N*_*cr*_ + *N*_*d*_ + *N*_*cn*_ = 435. The inclusion of these three groups (cancer types, cancer drugs, and countries) in the reduced set of *N*_*r*_ articles allows to investigate the interactions and influence of nodes inside group and between groups.

**Table 1.** List of articles devoted to cancer types in May 2017 English Wikipedia. This list of *N*_*cr*_ = 37 cancers taken from [24] is ordered by alphabetical order.

| Cancer type |  | Cancer type |  |
| --- | --- | --- | --- |
| 1 | Adrenal tumor | 21 | Mesothelioma |
| 2 | Anal cancer | 22 | Multiple myeloma |
| 3 | Appendix cancer | 23 | Neuroendocrine tumor |
| 4 | Bladder cancer | 24 | Non-Hodgkin lymphoma |
| 5 | Bone tumor | 25 | Oral cancer |
| 6 | Brain tumor | 26 | Ovarian cancer |
| 7 | Breast cancer | 27 | Pancreatic cancer |
| 8 | Cervical cancer | 28 | Prostate cancer |
| 9 | Cholangiocarcinoma | 29 | Skin cancer |
| 10 | Colorectal cancer | 30 | Soft-tissue sarcoma |
| 11 | Esophageal cancer | 31 | Spinal tumor |
| 12 | Gallbladder cancer | 32 | Stomach cancer |
| 13 | Gestational trophoblastic disease | 33 | Testicular cancer |
| 14 | Head and neck cancer | 34 | Thyroid cancer |
| 15 | Hodgkin’s lymphoma | 35 | Uterine cancer |
| 16 | Kidney cancer | 36 | Vaginal cancer |
| 17 | Leukemia | 37 | Vulvar cancer |
| 18 | Liver cancer |  |  |
| 19 | Lung cancer |  |  |
| 20 | Melanoma |  |  |

**Table 2.** List of articles devoted to cancer drugs in May 2017 English Wikipedia. This list of *N*_*d*_ = 203 cancer drugs taken from [25] is ordered by alphabetical order.

| Cancer drug | Cancer drug | Cancer drug | Cancer drug |
| --- | --- | --- | --- |
| 1 Abemaciclib | 52 Dactinomycin | 103 Ixazomib | 154 Prednisone |
| 2 Abiraterone acetate | 53 Daratumumab | 104 Lanreotide | 155 Procarbazine |
| 3 Acalabrutinib | 54 Dasatinib | 105 Lapatinib | 156 Propranolol |
| 4 Afatinib | 55 Daunorubicin | 106 Lenalidomide | 157 Protein-bound paclitaxel |
| 5 Aflibercept | 56 Decitabine | 107 Lenvatinib | 158 Radium-223 |
| 6 Alectinib | 57 Defibrotide | 108 Letrozole | 159 Raloxifene |
| 7 Alemtuzumab | 58 Degarelix | 109 Leuprorelin | 160 Ramucirumab |
| 8 Amifostine | 59 Denileukin diftitox | 110 Lomustine | 161 Rasburicase |
| 9 Aminolevulinic acid | 60 Denosumab | 111 Megestrol acetate | 162 Regorafenib |
| 10 Anastrozole | 61 Dexamethasone | 112 Melphalan | 163 Ribociclib |
| 11 Apalutamide | 62 Dexrazoxane | 113 Mercaptopurine | 164 Rituximab |
| 12 Aprepitant | 63 Dinutuximab | 114 Mesna | 165 Rolapitant |
| 13 Arsenic trioxide | 64 Docetaxel | 115 Methotrexate | 166 Romidepsin |
| 14 Asparaginase | 65 Doxorubicin | 116 Methylnaltrexone | 167 Romiplostim |
| 15 Atezolizumab | 66 Durvalumab | 117 Midostaurin | 168 Rucaparib |
| 16 Avelumab | 67 Elotuzumab | 118 Mitomycin C | 169 Ruxolitinib |
| 17 Axicabtagene ciloleucel | 68 Eltrombopag | 119 Mitoxantrone | 170 Siltuximab |
| 18 Axitinib | 69 Enzalutamide | 120 Necitumumab | 171 Sipuleucel-T |
| 19 Azacitidine | 70 Epirubicin | 121 Nelarabine | 172 Sonidegib |
| 20 Belinostat | 71 Eribulin | 122 Neratinib | 173 Sorafenib |
| 21 Bendamustine | 72 Erlotinib | 123 Netupitant/palonosetron | 174 Sunitinib |
| 22 Bevacizumab | 73 Etoposide | 124 Nilotinib | 175 Talc |
| 23 Bexarotene | 74 Everolimus | 125 Nilutamide | 176 Talimogene laherparepvec |
| 24 Bicalutamide | 75 Exemestane | 126 Niraparib | 177 Tamoxifen |
| 25 Bleomycin | 76 Filgrastim | 127 Nivolumab | 178 Temozolomide |
| 26 Blinatumomab | 77 Fludarabine | 128 Obinutuzumab | 179 Temsirolimus |
| 27 Bortezomib | 78 Fluorouracil | 129 Ofatumumab | 180 Thalidomide |
| 28 Bosutinib | 79 Flutamide | 130 Olaparib | 181 ThioTEPA |
| 29 Brentuximab vedotin | 80 Folinic acid | 131 Olaratumab | 182 Tioguanine |
| 30 Brigatinib | 81 Fulvestrant | 132 Omacetaxine mepesuccinate | 183 Tipiracil |
| 31 Busulfan | 82 Gefitinib | 133 Ondansetron | 184 Tisagenlecleucel |
| 32 Cabazitaxel | 83 Gemcitabine | 134 Osimertinib | 185 Tocilizumab |
| 33 Cabozantinib | 84 Gemtuzumab ozogamicin | 135 Oxaliplatin | 186 Topotecan |
| 34 Capecitabine | 85 Glucarpidase | 136 Paclitaxel | 187 Toremifene |
| 35 Carboplatin | 86 Goserelin | 137 Palbociclib | 188 Trabectedin |
| 36 Carfilzomib | 87 HPV vaccines | 138 Palifermin | 189 Trametinib |
| 37 Carmustine | 88 Hyaluronidase | 139 Palonosetron | 190 Trastuzumab |
| 38 Ceritinib | 89 Hydroxycarbamide | 140 Pamidronic acid | 191 Trastuzumab emtansine |
| 39 Cetuximab | 90 Ibritumomab tiuxetan | 141 Panitumumab | 192 Trifluridine |
| 40 Chlorambucil | 91 Ibrutinib | 142 Panobinostat | 193 Uridine triacetate |
| 41 Chlormethine | 92 Idarubicin | 143 Pazopanib | 194 Valrubicin |
| 42 Cisplatin | 93 Idelalisib | 144 Pegaspargase | 195 Vandetanib |
| 43 Cladribine | 94 Ifosfamide | 145 Pegfilgrastim | 196 Vemurafenib |
| 44 Clofarabine | 95 Imatinib | 146 Peginterferon | 197 Venetoclax |
| 45 Cobimetinib | 96 Imiquimod | 147 Pembrolizumab | 198 Vinblastine |
| 46 Copanlisib | 97 Inotuzumab ozogamicin | 148 Pemetrexed | 199 Vincristine |
| 47 Crizotinib | 98 Interferon alfa-2b | 149 Pertuzumab | 200 Vinorelbine |
| 48 Cyclophosphamide | 99 Interleukin 2 | 150 Plerixafor | 201 Vismodegib |
| 49 Cytarabine | 100 Ipilimumab | 151 Pomalidomide | 202 Vorinostat |
| 50 Dabrafenib | 101 Irinotecan | 152 Ponatinib | 203 Zoledronic acid |
| 51 Dacarbazine | 102 Ixabepilone | 153 Pralatrexate |  |

The paper is composed as follows: the section “Description of data sets and methods” will present the May 2017 English Wikipedia network, introduce the Google matrix, the PageRank and CheiRank algorithms, and explain the construction of reduced Google matrices. In this section the node influence is defined through the PageRank ranking and the PageRank sensitivity. In the section “Results” we present the influence of cancer devoted pages in Wikipedia and extract a cancer ranking which is compared to cancer rankings extracted from GBD study [5] and GLOBOCAN [2] databases. We also use the reduced Google matrix to construct a reduced network of cancers and we determine the interaction of cancers with countries and cancer drugs. We corroborate the results obtained from the network structure of Wikipedia articles with various disease burden and/or epidemiological studies. To our knowledge, it is the first time that such correspondence is established. Finally we compare cancer prescriptions obtained from May 2017 English Wikipedia network analysis with approved medications reported in National Cancer Institute [25] and DrugBank [29]. The last section presents the conclusion of this research.

## Description of data sets and methods

### Network of English Wikipedia articles of 2017

We analyze the English language edition of Wikipedia collected in May 2017 (ENWIKI2017) [30] containing *N* = 5 416 537 articles (nodes) connected by *N*_*l*_ = 122 232 932 directed hyperlinks between articles (without self-citations). From this data set we extract the *N*_*cr*_ = 37 types of cancers listed at [24]. From [25] we also collect names of drugs related to cancer diseases obtaining the list of *N*_*d*_ = 203 drugs present at Wikipedia. The lists of 37 cancer types and 203 drugs are given in Tabs. 1 and 2. This reduced set of *N*_*r*_ = 240 nodes is illustrated in the inset of Fig. 1. For global influence investigations, it is complemented by *N*_*cn*_ = 195 world countries listed in [28]. Thus in total we have the reduced network of *N*_*r*_ = *N*_*cr*_ + *N*_*d*_ + *N*_*cn*_ = 435 ≪ *N* nodes embedded in the global network with more than 5 millions nodes. All data sets are available at [28].

**Fig 1.**
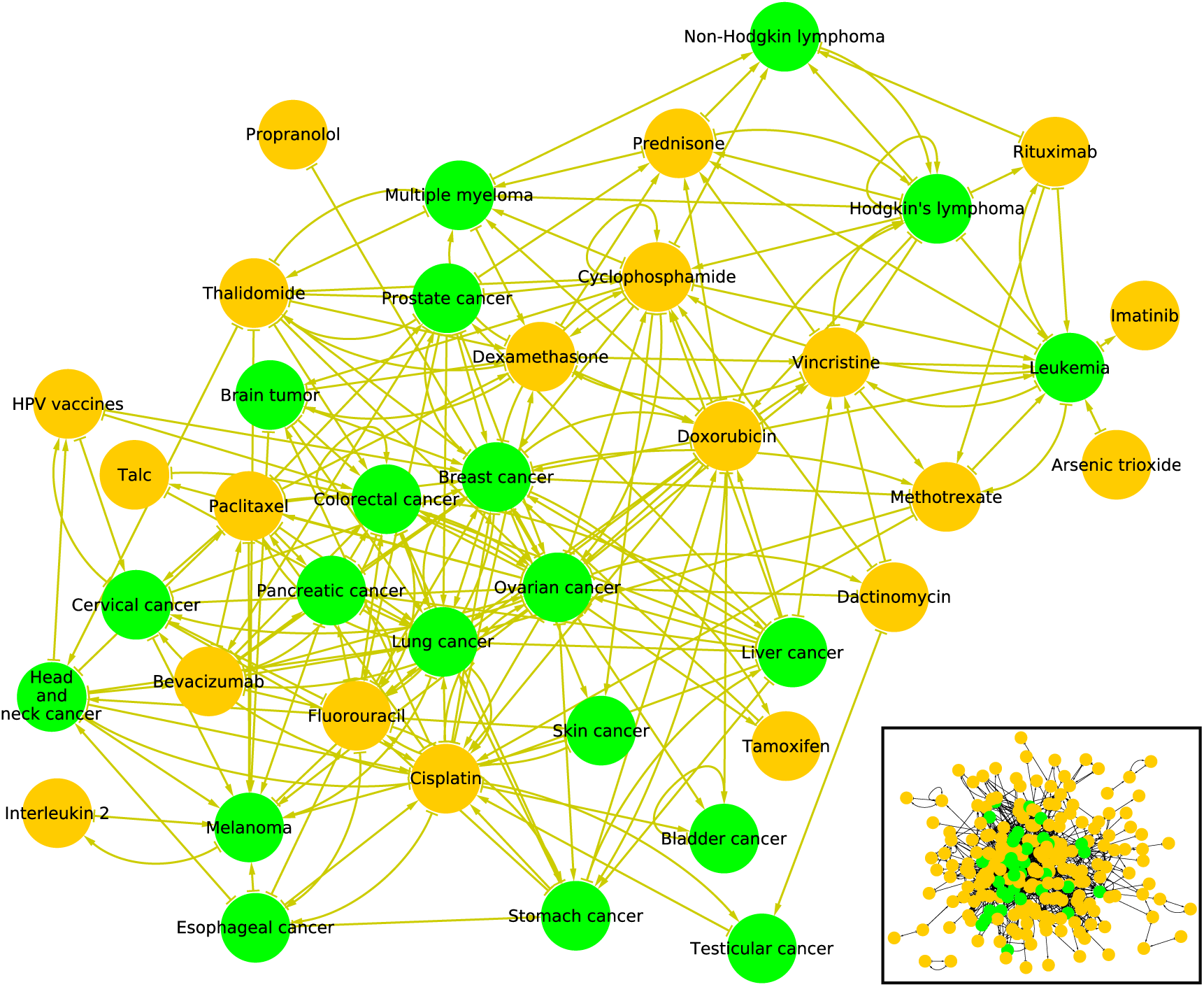
Subnetworks of cancers and cancer drugs in May 2017 English Wikipedia. Bottom right inset: subnetwork of *N*_*r*_ = 240 articles comprising *N*_*cr*_ = 37 articles devoted to cancers (green nodes) and *N*_*d*_ = 203 articles devoted to cancer drugs (golden nodes). Main figure: subnetwork of top 20 cancers and top 20 cancer drugs extracted from the ranking of 2017 English Wikipedia using PageRank algorithm (see Tab. 3). The bulk of the other Wikipedia articles is not shown. Arrows symbolize hyperlinks between cancer and cancer drug articles in the global Wikipedia.

### Google matrix construction rules

The construction rules of Google matrix *G* are described in detail in [6–8]. Thus the Google matrix *G* is built from the adjacency matrix *A*_*ij*_ with elements 1 if article (node) *j* points to article (node) *i* and zero otherwise. The Google matrix elements have the standard form *G*_*ij*_ = *αS*_*ij*_ + (1 − *α*)*/N* [6–8], where *S* is the matrix of Markov transitions with elements *S*_*ij*_ = *A*_*ij*_*/k*_*out*_(*j*). Here 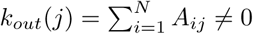 is the out-degree of node *j* (number of outgoing links) and *S*_*ij*_ = 1*/N* if *j* has no outgoing links (dangling node). The parameter 0 < *α* < 1 is the damping factor. For a random surfer, jumping from one node to another, it gives the probability (1 − *α*) to jump to any node. Below we use the standard value *α* = 0.85 [7] noting that for the range 0.5 ≤ *α* ≤ 0.95 the results are not sensitive to *α* [7, 8].

The right PageRank eigenvector of *G* is the solution of the equation *GP* = *λP* with the unit eigenvalue *λ* = 1. The PageRank components *P* (*j*) give positive probabilities to find a random surfer on a node *j* after an infinite journey (Σ_*j*_ *P* (*j*) = 1). The numerical computation of *P* (*j*) is done efficiently with the PageRank algorithm described in [6, 7].

The node influence is measured from the PageRank algorithm. We sort network nodes by decreasing PageRank probabilities. We assign *K* = 1 index to the node with maximal probability, i.e., the most central node according to PageRank algorithm, *K* = 2 index to the node with the second biggest probability, … A recursive definition of the PageRank algorithm can be given: a node is all the more influential as it is pointed by influential nodes.

It is also useful to consider the network with inverted direction of links. After links inversion 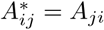, the Google matrix *G*\* is constructed within the same procedure with *G*\**P*\* = *P*\*. The matrix *G*\* has its own PageRank vector *P*\* called CheiRank [31] (see also [8, 32]). Its probability values can be again ordered in a decreasing order with CheiRank index *K*\* with highest *P*\*(*j*) at *K*\* = 1 and smallest at *K*\* = *N*. The CheiRank algorithm measures the node diffusivity. A recursive definition of the CheiRank algorithm can also be given: a node is all the more diffusive as it is pointed by diffusive nodes.

On average, the high values of *P* (*j*) (*P*\*(*j*)) correspond to nodes *j* with many ingoing (outgoing) links [8].

The PageRank order list of 37 cancers and 203 drugs is given in Table 3. In the global ENWIKI2017 network, countries are located on top PageRank positions (1. *USA*, 4. *France*, 5. *Germany*) so that cancers and drugs are located well below them since the first cancer type, i.e. *Lung cancer*, appears at 3 478th position, and the first cancer drug, i.e. *Talc*, appears at 22 177th position (see Fig. 2). As expected cancer types have a more central position than cancer drugs. The network of 40 nodes and their direct links is shown in Fig. 1 for the top 20 PageRank nodes of cancers and drugs (ordered separately for cancers and drugs). We see that already only for 40 nodes the network structure is rather complex. Here and below the networks are drawn with Cytoscape [33].

**Table 3.**
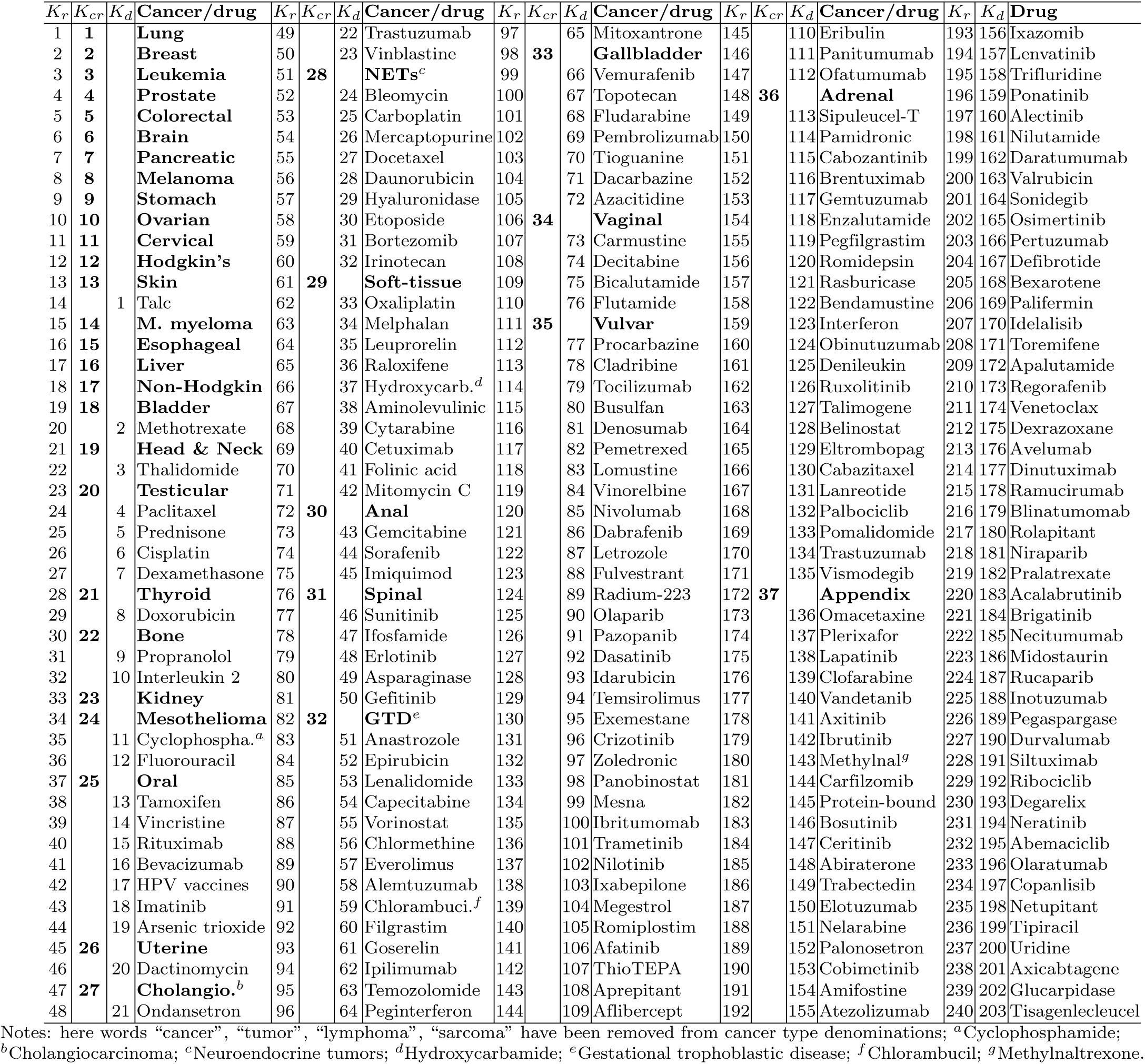
Ranking of articles devoted to cancer types and to cancer drugs in May 2017 English Wikipedia using PageRank algorithm. Cancer types are highlighted in boldface.

**Fig 2.**
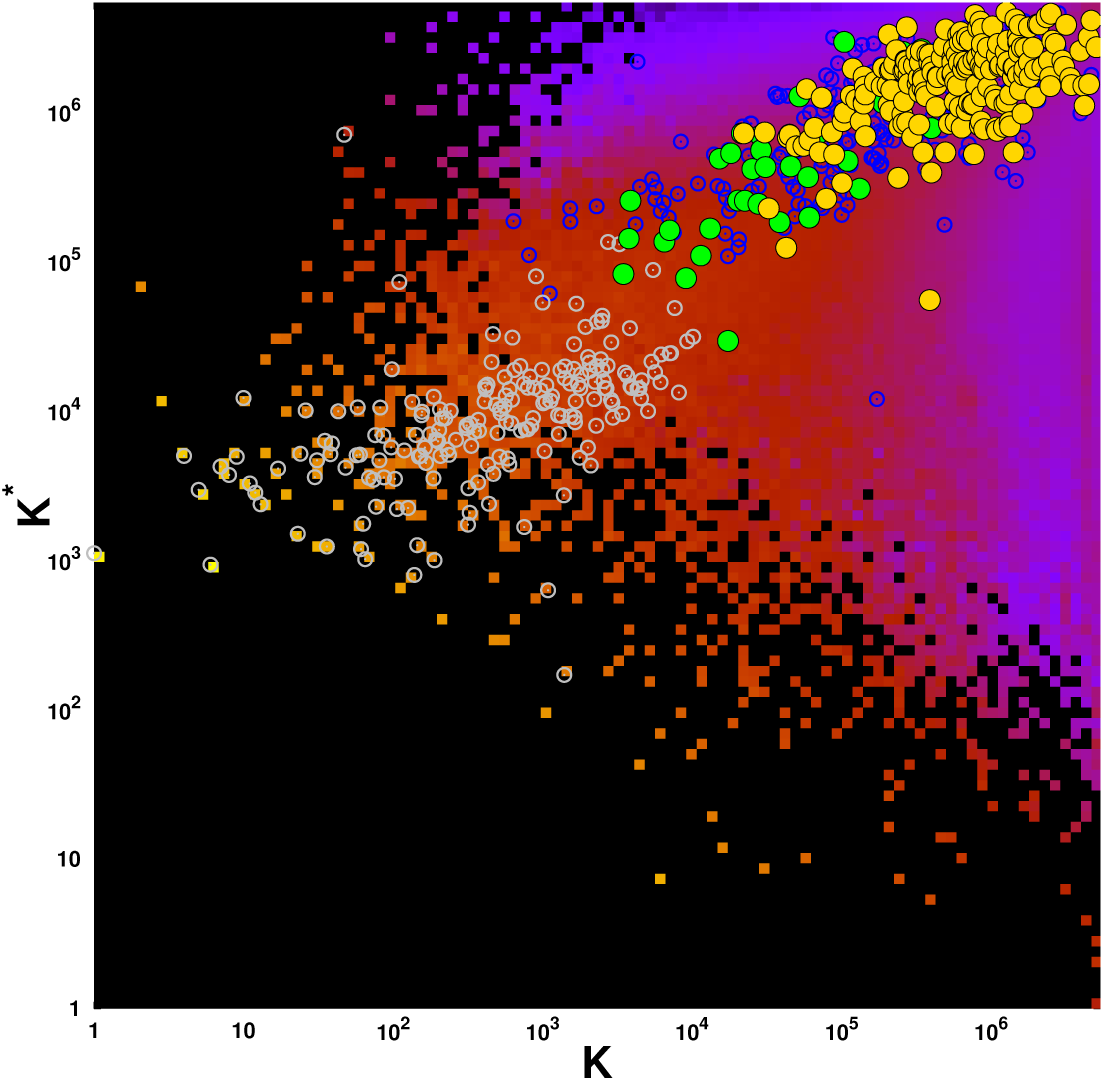
Density of May 2017 English Wikipedia articles in the CheiRank *K*\* – PageRank *K* plane. Data are averaged over a 100 × 100 grid spanning the (log_10_ *K*, log_10_ *K*\*) ∈ [0, log_10_ *N*] × [0, log_10_ *N*] domain. Density of articles ranges from very low density (purple tiles) to very high density (bright yellow tiles). The absence of article is represented by black tiles. The superimposed green (gold) circles give the positions of May 2017 English Wikipedia articles devoted to cancers (cancer drugs) listed in Tab. 1 (Tab. 2). For comparison, the gray (blue) open circles give the positions of pages devoted to sovereign countries (infectious diseases) in May 2017 English Wikipedia.

### Reduced Google matrix algorithm

The details of REGOMAX method are described in [16, 17, 20]. It captures in the reduced Google matrix of size *N*_*r*_ × *N*_*r*_ the full contribution of direct and indirect pathways existing in the full Google matrix between *N*_*r*_ nodes of interest. The reduced Google matrix *G*_R_ is such as *G*_R_*P*_*r*_ = *P*_*r*_ where *P*_*r*_ is its associated PageRank probability vector. The PageRank probabilities *P*_*r*_(*j*) of the selected *N*_*r*_ nodes are the same as for the global network with *N* nodes, up to a constant multiplicative factor taking into account that the sum of PageRank probabilities over *N*_*r*_ nodes is unity. The computation of *G*_R_ provides a decomposition into matrices that clearly distinguish direct from indirect interactions: *G*_R_ = *G*_*rr*_ + *G*_pr_ + *G*_qr_ [17]. Here *G*_*rr*_ is the *N*_*r*_ × *N*_*r*_ submatrix of the *N* × *N* global Google matrix *G* encoding the direct links between the selected *N*_*r*_ nodes. The *G*_pr_ matrix is rather close to the matrix in which each column is given by the PageRank vector *P*_*r*_, ensuring that PageRank probabilities of *G*_R_ are the same as for *G* (up to a constant multiplier). Thus *G*_pr_ does not provide much more information about direct and indirect links between selected nodes than the usual Google matrix analysis described in the previous section. The component playing an interesting role is *G*_qr_, which takes into account all indirect links between selected nodes appearing due to multiple paths via the global network of *N* nodes (see [16, 17]). The matrix *G*_qr_ = *G*_qrd_ + *G*_qrnd_ has diagonal (*G*_qrd_) and non-diagonal (*G*_qrnd_) parts. Thus *G*_qrnd_ describes indirect interactions between nodes. The explicit formulas as well as the mathematical and numerical computation methods of all three components of *G*_R_ are given in [16, 17, 20].

With the reduced Google matrix *G*_R_ and its components we can analyze the PageRank sensitivity in respect to specific links between *N*_*r*_ nodes. To measure the sensitivity of a country *cn* to a cancer *cr* we change the matrix element (*G*_R_)_*cn,cr*_ by a factor (1 + *δ*) with *δ* ≪ 1 and renormalize to unity the sum of the column elements associated with cancer *cr*, and we compute the logarithmic derivative of PageRank probability *P* (*cn*) associated to country *cn*: *D*(*cr* → *cn, cn*) = *d* ln *P* (*cn*)*/dδ* (diagonal sensitivity). It is also possible to consider the nondiagonal (or indirect) sensitivity *D*(*cr* → *cn, cn*’) = *d* ln *P* (*cn*’)*/dδ* when the variation is done for the link from *cr* to *cn* and the derivative of PageRank probability is computed for another country *cn*’. Also instead of the link *cr* → *cn* we can consider the link from a cancer *cr* to a drug *d* computing then the nondiagonal sensitivity of country *cn*’. The PageRank sensitivity approach, already used in [26, 27], allows to measure the sensitivity of a node influence with respect to the change of a link intensity. Pragmatically it measures the increase or decrease of influence of a node *A* caused by an increase of the intensity of the link from a node *B* to a node *C*.

## Results

### Cancer distribution on PageRank-CheiRank plane

The PageRank order of 37 cancers and 203 cancer drugs is given in Tab. 3. In the top 3 positions we find *Lung, Breast, Leukemia* cancers. *Lung* and *Breast cancers* have indeed the two highest incidences [2] and *Leukemia* is the most frequent type of cancer in children and young adults [34]. In general in the PageRank order of 240 cancers and drugs, cancers occupy predominantly the top positions. The first three drugs are *Talc, Methotrexate, Thalidomide*, taking positions 14, 20, 22. The top position of *Talc* among cancer drugs may be explained by its industrial use and also by both potential carcinogenic and anticancer effects [35]. *Methotrexate* can be used in the most frequent types of cancer but also in autoimmune diseases and for medical abortions [36]. The third position of *Thalidomide* among cancer drugs may be explained by its high potential for the treatment of cancers but also for its well-known teratogenic effect; this teratogenic effect may by itself contribute to its prominence in Wikipedia. It is also used for treatment of other diseases than cancers (tuberculosis, graft-versus-host disease,…) [37]. The list of these 240 articles in CheiRank order is also given in [28].

The distribution of selected articles on the global PageRank-CheiRank plane of the whole Wikipedia network with *N* = 5 416 537 nodes are shown in Fig. 2. The top PageRank positions are taking by the world countries as discussed in [8, 26] marked by gray open circles. Then there is a group of cancers (above *K* ∼ 3 × 10^3^ and *K*\* ∼ 10^4^), marked by green points, followed by drugs (mostly above *K* ∼ 10^4^ and *K*\* ∼ 10^5^), marked by gold points. There is a certain overlap between cancers and drugs on this plane but in global there is a clear separation between these two groups. As a comparison we also mark the positions of 230 infectious diseases by open blue circles. These 230 articles are studied in [27] in the frame of Wikipedia network analysis. The global PageRank list of 230 infectious diseases and 37 cancers is given in [28]. In this list *Lung cancer* is located at the 7th position. From Fig. 2 we observe these two types of diseases occupy somewhat the same (*K, K*\*) region (mostly above *K*\* ∼ 10^5^ and above *K* ∼ 3 × 10^3^) suggesting that cancer types and infectious diseases have globally the same influence in May 2017 English Wikipedia with the exception of the first six infectious diseases, *Tuberculosis* (*K* = 639), *HIV/AIDS* (*K* = 810), *Malaria* (*K* = 1116), *Pneumonia* (*K* = 1531), *Smallpox* (*K* = 1532), *Cholera* (*K* = 2300) which are causing or have caused pandemics or notable epidemics. The first three cancer types, i.e. *Lung cancer, Breast cancer*, and *Leukemia*, appear at positions *K* = 3478, 3788, and 3871 just before *Influenza* at *K* = 4191.

The 240 cancer types and drugs placed on the plane of local PageRank indices *K*_*r*_ ∈ {1, …, 240} and CheiRank indices 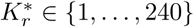 is shown in Fig. 3. We retrieve the fact that cancer types occupy the top positions in *K*_*r*_ and in 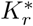. Indeed the first 14 most influent articles of this subset (*K* ≤ 14), which appear to be devoted to cancer types, are also the most communicative with the exception of articles devoted to drugs *Paclitaxel* 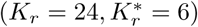 and *Bicalutamide* 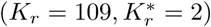. *Paclitaxel* [38] is a chemotherapy medication used to treat a wide range of cancer types e.g. *Ovarian cancer, Breast cancer, Lung cancer, Pancreatic cancer*, etc. Moreover *Paclitaxel* article cites *Ovarian cancer* article 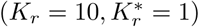 which is a very communicative article since the *Ovarian cancer* article CheiRank index, *K*\* = 29 317, is about one order magnitude lower than the CheiRank indexes, *K*\* ≳ 10^5^, of the other 239 considered articles (see Fig. 2). The wide applications of *Paclitaxel* and the citation of *Ovarian cancer* article explain the very good ranking of this cancer drug in the CheiRank scale. On the other hand, the 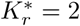 rank of the *Bicalutamide* article (see Fig. 3), devoted to an antiandrogen medication mainly used to treat *Prostate cancer*, is due to a very long article with a high density of intra-wiki citations [39]. Like the *Paclitaxel* article, the *Bicalutamide* article cites also the *Ovarian cancer* since this medication has already been tried for this cancer type [39].

**Fig 3.**
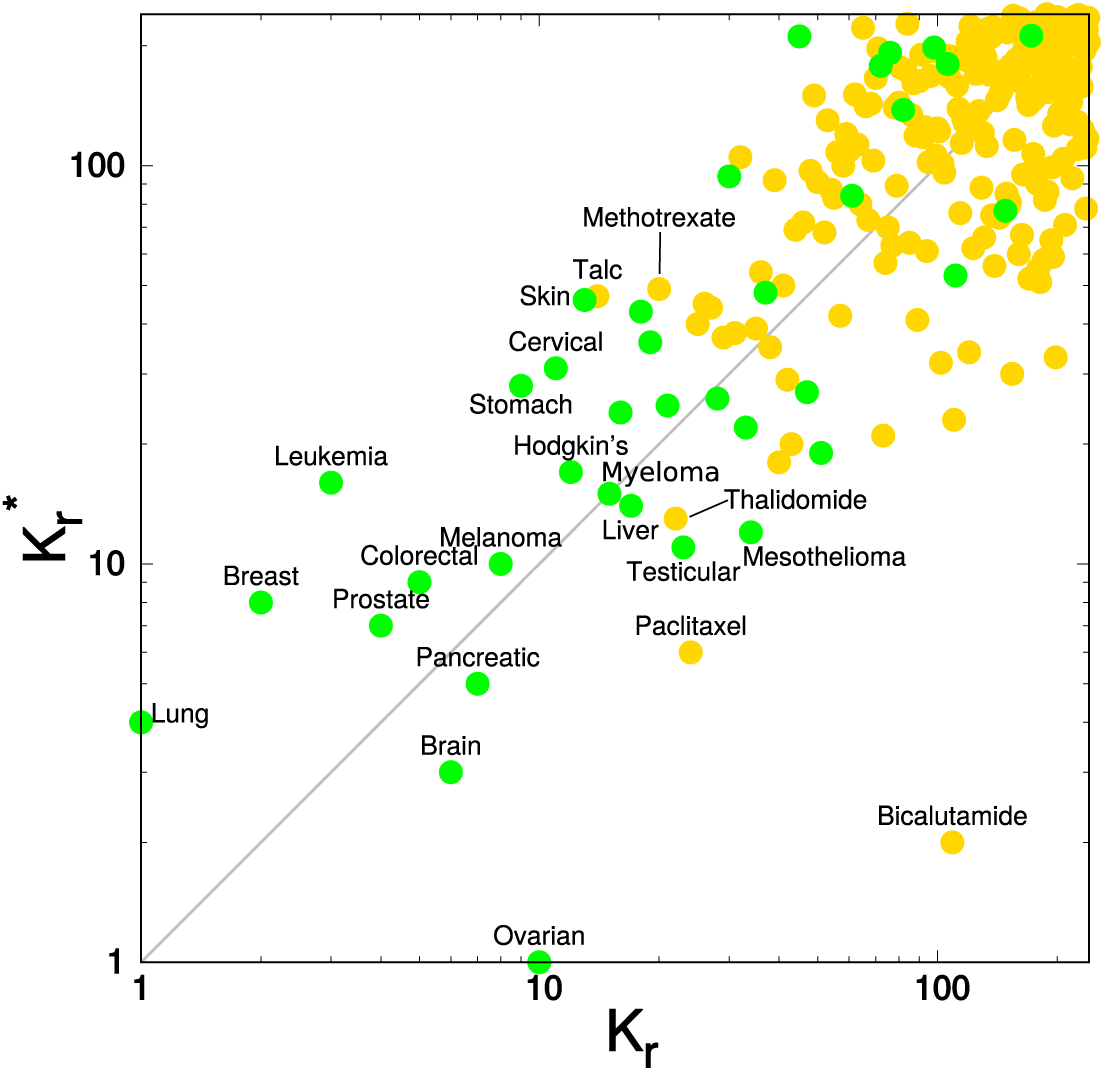
Distribution of the May 2017 English Wikipedia articles devoted to cancers and drug cancers in the local CheiRank 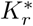 — PageRank *K*_*r*_ plane. The *N*_*cr*_ = 37 (*N*_*d*_ = 203) articles devoted to cancers (drug cancers) are represented by green (gold) plain circles.

The three most influent cancer drugs in ENWIKI2017 are *Talc, K*_*r*_ = 14, which is used to prevent blood effusions, e.g., in *Lung cancer* or *Ovarian cancer* [35], *Methotrexate, K*_*r*_ = 20, which is a chemotherapy agent used for the treatment *Breast cancer, Leukemia, Lung cancer, Lymphoma*, etc [36], and *Thalidomide, K*_*r*_ = 22, which is a drug modulating the immune system used, e.g., for *Multiple myeloma* treatment [37]. Although *Talc* is widely used in chemical, pharmaceutical and food industries [35], its global PageRank position is nevertheless of the same order than the PageRank position of the second most influent cancer drug in Wikipedia, i.e., *Methotrexate*, which is a drug more specific to cancers [36].

### Comparison of Wikipedia network analysis with GBD study 2017 and GLOBOCAN 2018 for cancer significance

We perform the comparison of cancer significance given by the GBD study 2017 [5], the GLOBOCAN 2018 [2], and the Wikipedia network analysis. We extract the rankings of cancer types by the number of deaths in 2017 estimated by the 2017 GBD study [40] (see Tab. 4) and by the number of disability-adjusted life years (DALYs) estimated by the 2017 GBD study [41] (see Tab. 4). Also, we extract the rankings of cancer types by the number of deaths and by the number of new cases in 2018 estimated by the GLOBOCAN 2018 [4] (see Tab. 5). In Fig. 4, we show the overlap of these 4 rankings with the extracted ranking of cancer types obtained from the ENWIKI2017 PageRanking (see bold items in Tab. 3). We observe that the ranking obtained from the Wikipedia network analysis provides a reliable cancer types ranking since its top 10 (top 20) shares about 70% (80%) similarity with GBD study data and GLOBOCAN data. The Wikipedia top 5 reaches even 80% similarity with top 5 cancer types extracted from the estimated number of new cases in 2018.

**Fig 4.**
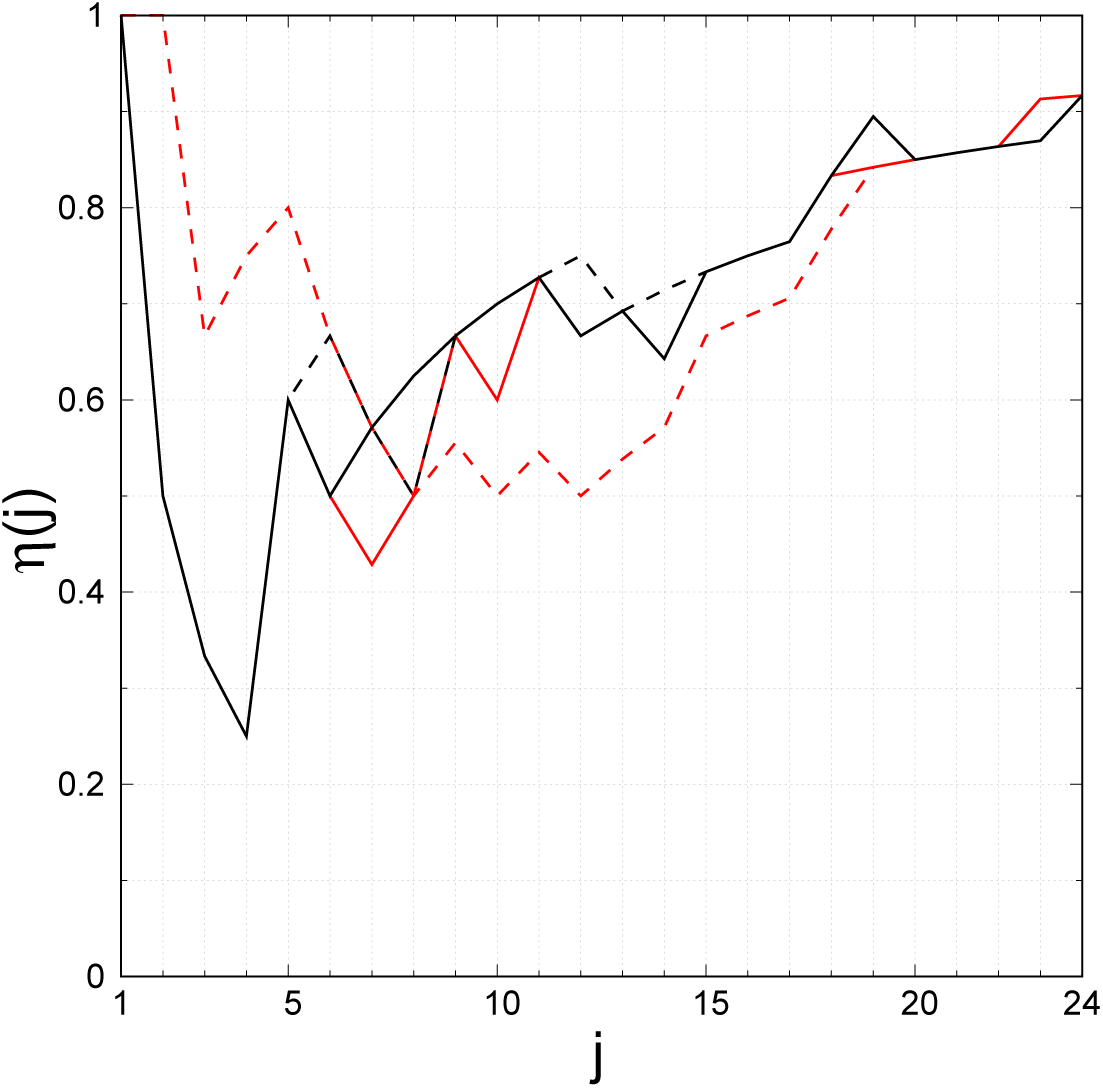
Comparison between cancer rankings extracted from May 2017 English Wikipedia PageRank, from the global burden of disease (GBD) study 2017 data, and from GLOBOCAN 2018 data. The overlap *η*(*j*) gives the number of cancer types in common in the top *j* of the ranking of cancers obtained from the May 2017 English Wikipedia PageRank (see bold terms in Tab. 3) and in the top *j* of the ranking of cancers by estimated number of worldwide deaths from GBD 2017 data [40] (black line, see Tab. 4), by estimation of disability-adjusted life years from GBD 2017 data [41] (black dashed line, Tab. 4), by estimated number of worldwide deaths from GLOBOCAN 2018 data [4] (red line, Tab. 5), and by estimated number of new cases from GLOBOCAN 2018 data [4] (red dashed line, Tab. 5). Only the black plain line is visible, where black plain line, red plain line and black dashed line overlap, e.g., from *j* = 1 to *j* = 5.

**Table 4.** List of cancer types ordered by the estimated number of deaths during the year 2017 (left table) and by the estimated disability-adjusted life years (DALYs) for 2017 (right table). Data extracted from GBD Study [40, 41].

| Rank | Cancer | Deaths<br>in 2017 ($\times 10^3$) | Rank | Cancer | DALYs<br>in 2017 ($\times 10^3$) |
| --- | --- | --- | --- | --- | --- |
| 1 | Lung cancer | 1883.1 | 1 | Lung cancer | 40900 |
| 2 | Colorectal cancer | 896.0 | 2 | Liver cancer | 20800 |
| 3 | Stomach cancer | 865.0 | 3 | Stomach cancer | 19100 |
| 4 | Liver cancer | 819.4 | 4 | Colorectal cancer | 19000 |
| 5 | Breast cancer | 611.6 | 5 | Breast cancer | 17700 |
| 6 | Pancreatic cancer | 441.1 | 6 | Leukemia | 12000 |
| 7 | Esophageal cancer | 436.0 | 7 | Head and neck cancer | 10600 |
| 8 | Prostate cancer | 415.9 | 8 | Esophageal cancer | 9780 |
| 9 | Head and neck cancer | 380.6 | 9 | Pancreatic cancer | 9080 |
| 10 | Leukemia | 347.6 | 10 | Brain tumor | 8740 |
| 11 | Cervical cancer | 259.7 | 11 | Cervical cancer | 8060 |
| 12 | Non-Hodgkin lymphoma | 248.6 | 12 | Prostate cancer | 7060 |
| 13 | Brain tumor | 247.1 | 13 | Non-Hodgkin lymphoma | 7020 |
| 14 | Bladder cancer | 196.5 | 14 | Ovarian cancer | 4670 |
| 15 | Ovarian cancer | 176.0 | 15 | Bladder cancer | 3600 |
| 16 | Gallbladder cancer | 174.0 | 16 | Gallbladder cancer | 3480 |
| 17 | Kidney cancer | 138.5 | 17 | Kidney cancer | 3280 |
| 18 | Skin cancer | 126.8 | 18 | Skin cancer | 2980 |
| 19 | Multiple myeloma | 107.1 | 19 | Multiple myeloma | 2330 |
| 20 | Uterine cancer | 85.2 | 20 | Uterine cancer | 2140 |
| 21 | Thyroid cancer | 41.2 | 21 | Hodgkin's lymphoma | 1380 |
| 22 | Hodgkin's lymphoma | 32.6 | 22 | Thyroid cancer | 1130 |
| 23 | Mesothelioma | 29.9 | 23 | Mesothelioma | 671 |
| 24 | Testicular cancer | 7.7 | 24 | Testicular cancer | 375 |

**Table 5.**
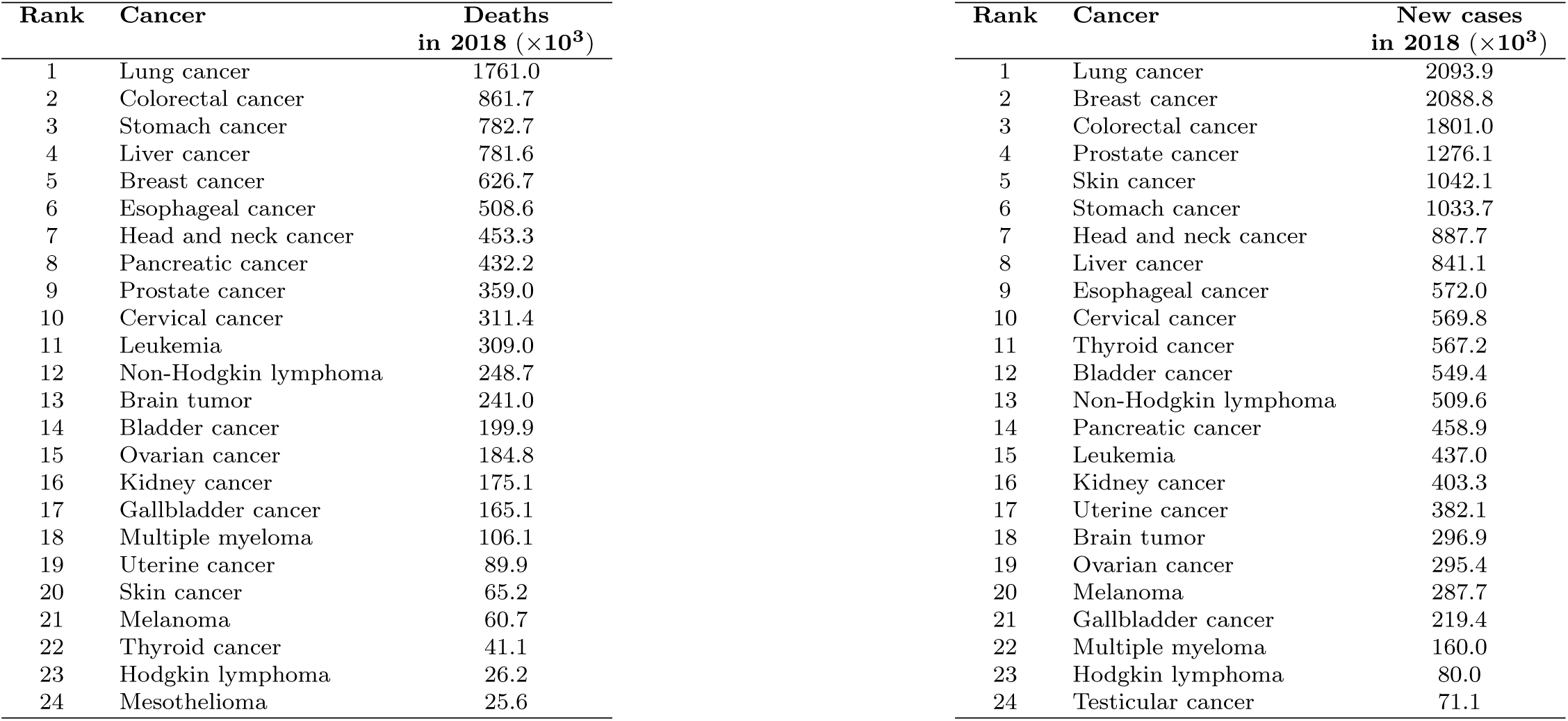
List of cancer types ordered by the estimated number of deaths during the year 2018 (left table) and by the estimated number of new cases in 2018 (right table). Data extracted from GLOBOCAN 2018 [4]

| Rank | Cancer | Deaths<br>in 2018 ($\times 10^3$) | Rank | Cancer | New cases<br>in 2018 ($\times 10^3$) |
| --- | --- | --- | --- | --- | --- |
| 1 | Lung cancer | 1761.0 | 1 | Lung cancer | 2093.9 |
| 2 | Colorectal cancer | 861.7 | 2 | Breast cancer | 2088.8 |
| 3 | Stomach cancer | 782.7 | 3 | Colorectal cancer | 1801.0 |
| 4 | Liver cancer | 781.6 | 4 | Prostate cancer | 1276.1 |
| 5 | Breast cancer | 626.7 | 5 | Skin cancer | 1042.1 |
| 6 | Esophageal cancer | 508.6 | 6 | Stomach cancer | 1033.7 |
| 7 | Head and neck cancer | 453.3 | 7 | Head and neck cancer | 887.7 |
| 8 | Pancreatic cancer | 432.2 | 8 | Liver cancer | 841.1 |
| 9 | Prostate cancer | 359.0 | 9 | Esophageal cancer | 572.0 |
| 10 | Cervical cancer | 311.4 | 10 | Cervical cancer | 569.8 |
| 11 | Leukemia | 309.0 | 11 | Thyroid cancer | 567.2 |
| 12 | Non-Hodgkin lymphoma | 248.7 | 12 | Bladder cancer | 549.4 |
| 13 | Brain tumor | 241.0 | 13 | Non-Hodgkin lymphoma | 509.6 |
| 14 | Bladder cancer | 199.9 | 14 | Pancreatic cancer | 458.9 |
| 15 | Ovarian cancer | 184.8 | 15 | Leukemia | 437.0 |
| 16 | Kidney cancer | 175.1 | 16 | Kidney cancer | 403.3 |
| 17 | Gallbladder cancer | 165.1 | 17 | Uterine cancer | 382.1 |
| 18 | Multiple myeloma | 106.1 | 18 | Brain tumor | 296.9 |
| 19 | Uterine cancer | 89.9 | 19 | Ovarian cancer | 295.4 |
| 20 | Skin cancer | 65.2 | 20 | Melanoma | 287.7 |
| 21 | Melanoma | 60.7 | 21 | Gallbladder cancer | 219.4 |
| 22 | Thyroid cancer | 41.1 | 22 | Multiple myeloma | 160.0 |
| 23 | Hodgkin lymphoma | 26.2 | 23 | Hodgkin lymphoma | 80.0 |
| 24 | Mesothelioma | 25.6 | 24 | Testicular cancer | 71.1 |

### Reduced Google matrix of cancers and drugs

Let us consider now the subset of *N*_*r*_ = 40 nodes composed of the first 20 cancers and the first 20 cancer drugs of the ENWIKI2017 PageRanking (Tab. 3). For this sub-network of interest illustrated in Fig. 1, we perform the calculation of the reduced Google matrix *G*_R_ and its components *G*_*rr*_, *G*_pr_ and, *G*_qr_. From Fig. 5, as expected, we observe that the *G*_R_ matrix (top left panel) is dominated by the *G*_pr_ component (bottom left panel) since *W*_pr_ = 0.872*W*_R_. The *G*_pr_ component is of minor interest as it expresses again the relative PageRanking between the *N*_*r*_ = 40 cancers and drugs already obtained and discussed in previous sections. The *G*_*rr*_ (top right panel) gives the direct links between the considered cancers and drugs. Indeed, the *G*_*rr*_ matrix is similar to the adjacency matrix *A* since there is a one-to-one correspondence between non zero entries of *G*_*rr*_ and of *A* (for *G*_*rr*_ by non zero entry we mean an entry greater than (1 − *α*)*/N ≃* 2.8 × 10^−8^). Fig. 1 illustrates the subnetwork of the direct links between the top 20 cancer types and the top 20 cancer drugs encoded in *G*_*rr*_ and *A*. Once the obvious *G*_pr_ component and the direct links *G*_*rr*_ component removed from the reduced Google matrix *G*_R_, the remaining part *G*_qr_ gives the hidden links between the set of *N*_*r*_ nodes of interest. In Fig. 5 we represent *G*_qrnd_ (bottom right panel), the non diagonal part of *G*_qr_. We can consider that a link with a non zero entry in *G*_qrnd_ and a zero entry in *G*_*rr*_ (consequently also in *A*) is a hidden link. Below we use the non obvious components of *G*_*rr*_ + *G*_qrnd_ to draw the structure of reduced network.

**Fig 5.**
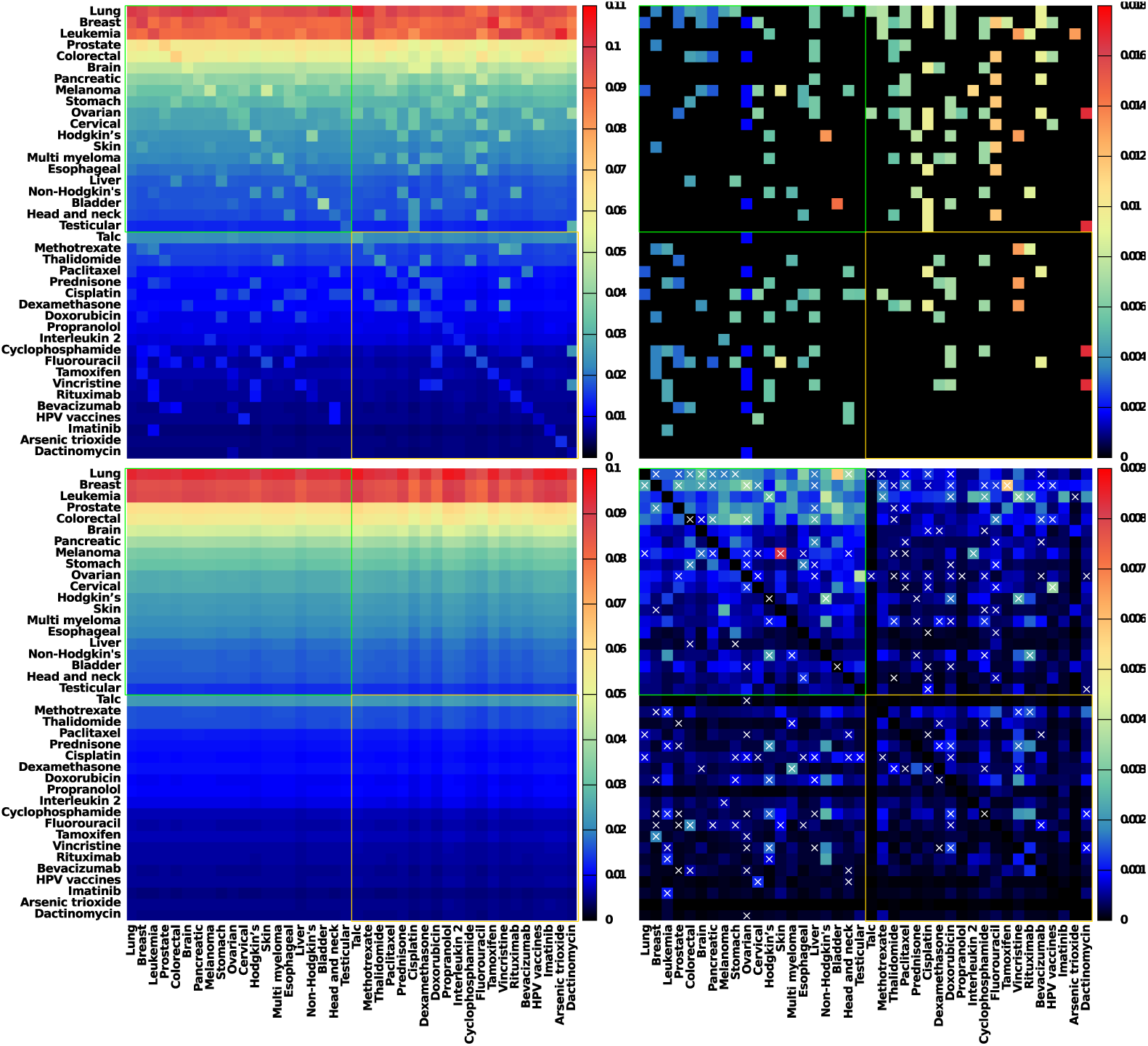
Reduced Google matrix *G*_R_ associated to the intertwined subnetworks of top 20 cancer articles and of top 20 drug articles. The reduced Google matrix *G*_R_ (top left) and its 3 components *G*_*rr*_ (top right), *G*_pr_ (bottom left), and *G*_qrnd_ (bottom right) are shown. The weights of the components are *W*_R_ = 1, *W*_pr_ = 0.872, *W*_*rr*_ = 0.086, and *W*_qr_ = 0.042 (*W*_qrnd_ = 0.038). For each component, thin green and gold lines delimit cancers and drugs sectors, i.e. upper left sub-matrix characterizes *from cancers to cancers* interactions, lower right sub-matrix *from drugs to drugs* interactions, upper right sub-matrix *from drugs to cancers* interactions, and lower left sub-matrix *from cancers to drugs* interactions. On the *G*_qrnd_ component (bottom right) superimposed crosses indicate links already present in the adjacency matrix (otherwise stated links corresponding to non zero entries in *G*_*rr*_, see top right).

### Reduced network of cancers

We construct the reduced Google matrix associated to the set of *N*_*r*_ = *N*_*cr*_ + *N*_*cn*_ = 232 Wikipedia articles constituted of *N*_*cr*_ = 37 articles devoted to cancer types and of *N*_*cn*_ = 195 articles devoted to countries. We consider the top 5 cancer types appearing in the ranking of May 2017 English Wikipedia using the PageRank algorithm which, according to Tab. 3, are 1 *Lung cancer*, 2 *Breast cancer*, 3 *Leukemia*, 4 *Prostate cancer*, 5 *Colorectal cancer*. Let us ordinate cancer types by their relative ranking in Tab. 3, cancer type *cr*_*i*_ is consequently the *i*th most influent cancer type in May 2017 English Wikipedia. Using the reduced Google matrix, the component (*G*_*rr*_ + *G*_qrnd_)_*cr*i_,*cr*_j_, where *i, j* ∈ {1, …, *N*_*cr*_}, gives the non obvious strength of the link pointing from the *j*th to the *i*th most influent cancer types. From each one the top 5 cancer types, {*cr*_*j*_}_*j*∈{1,…,5}_, we select the two cancer types 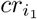 and 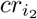, with *i*_1_, *i*_2_ ∈ {1, …, *j* − 1, *j* + 1, …, *N*_*cr*_}, to which cancer type *cr*_*j*_ is preferentially linked (“friends”), i.e. those giving the two strongest (*G*_*rr*_ + *G*_qrnd_)_*cr*i,*cr*j_ components. Around the main circle in Fig. 6 (top panel) we first place the top 5 most influent cancer types in May 2017 English Wikipedia. Then we connect each one of these cancer types to their two above defined cancer type friends. If these cancer types are not yet present in the network we add them in the vicinity of the cancer type pointing them. For each newly added cancer type we reiterate the same process until no new cancer type is added to the reduced network. The construction process of the reduced network of cancer ends at the 3rd iteration (see Fig. 6, top panel) exhibiting only 10 of the *N*_*cr*_ = 37 cancer types, which in addition of the top 5 cancer types, are 8 *Melanoma*, 9 *Stomach cancer*, 12 *Hodgkin lymphoma*, 17 *Liver cancer* and 18 *Non-Hodgkin lymphoma*. Among these 10 cancer types, 7 are among the top 10 deadliest in 2017 according to GBD study (see Tab. 4). In the reduced network of cancers showed in Fig. 6 (top panel) we observe that the most influent cancer, i.e., *Lung cancer* is pointed from all the other cancer types with the exception of *Hodgkin and Non-Hodgkin lymphomas*. Also, Fig. 6 (top panel) exhibits clearly a cluster of cancers (*Colorectal, Stomach*, and *Liver cancers*) affecting the digestive system, a cluster of cancers (*Hodgkin and Non-Hodgkin lymphomas*, and *Leukemia*) affecting blood, a loop interaction between *Prostate* and *Breast cancers* which are both linked to steroid hormone pathways and may be both treated with hormone therapy [42, 43], loop interactions between *Lung* and *Breast cancers* and between *Lung cancer* and *Melanoma* affecting mainly the thoracic region.

**Fig 6.**
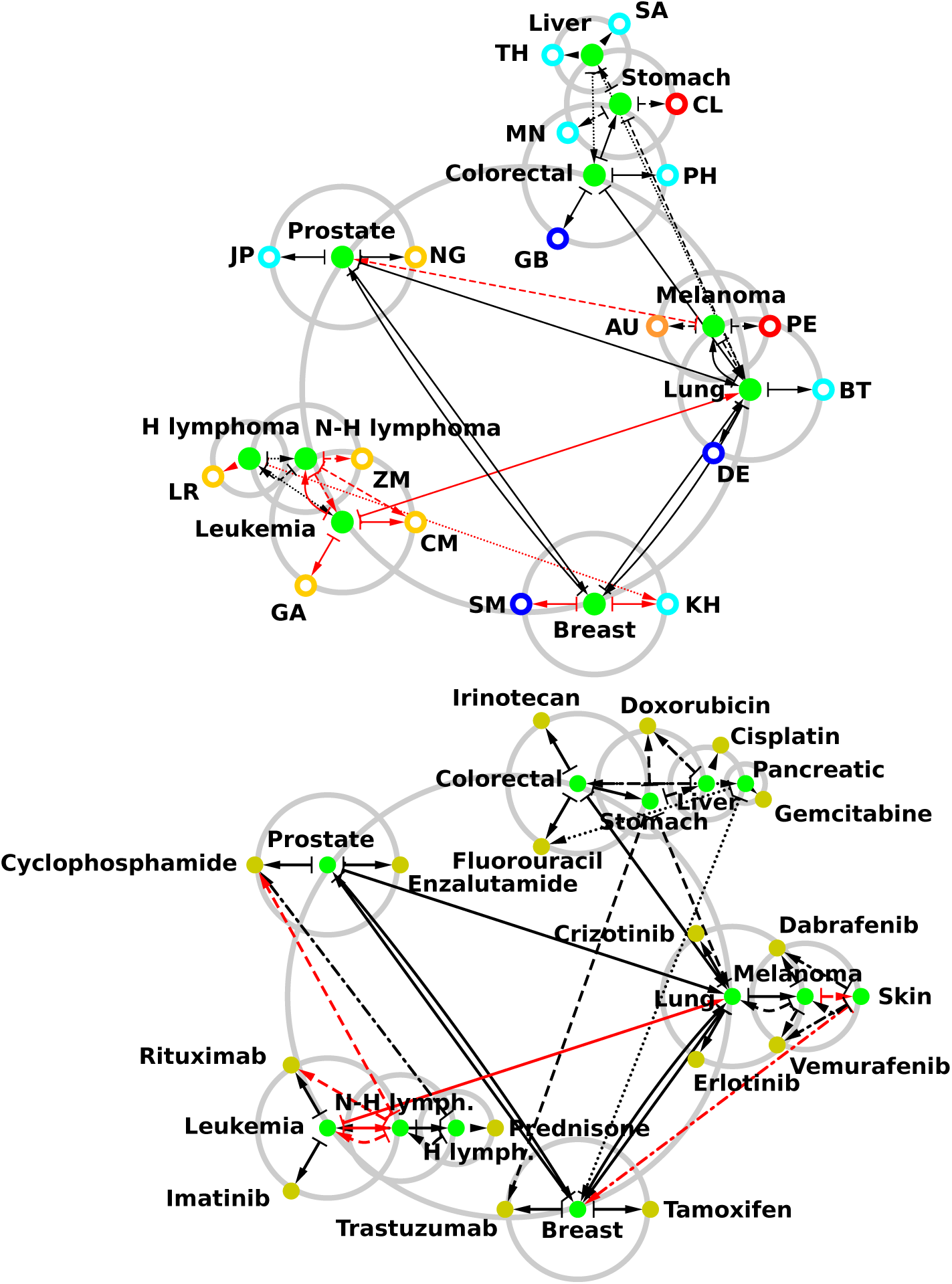
Reduced network of cancers. We consider the reduced Google matrix associated to the *N*_*cr*_ = 37 cancers and (top panel) the *N*_*cn*_ = 195 countries, (bottom panel) the *N*_*d*_ = 203 cancer drugs. We consider the top 5 cancers from the ranking of May 2017 English Wikipedia using the PageRank algorithm: 1. *Lung cancer*, 2. *Breast cancer*, 3. *Leukemia*, 4. *Prostate cancer*, 5. *Colorectal cancer* (see Tab. 3). These 5 cancers are symbolized by plain green nodes distributed around the central gray circle. We determine the two cancers to which each of these 5 cancers are preferentially linked according to (*G*_*rr*_ + *G*_qrnd_). If not among the top 5 cancers, a newly determined cancer is placed on a gray circle centered on the cancer from which it is linked. Then for each one of the newly added cancers we determine the two best cancers to which they are each linked, and so on. This process is stopped once no new cancers can be added, i.e. at the 3rd iteration (top panel) and 4th iteration (bottom panel). Also, at each iteration the two countries (drugs) to which each cancer are preferentially linked are placed on the gray circle centered on the cancer; see top panel (bottom panel). No new links are determined from the newly added countries or drugs. On top panel, countries are represented by ring shaped nodes (red for American countries, yellow for African countries, cyan for Asian countries, blue for European countries, and orange for Oceanian countries). On bottom panel, drugs are represented by plain gold nodes. The arrows represent the directed links between cancers and from cancers to countries or drugs (1st iteration: plain line; 2nd iteration: dashed line; 3rd iteration: dotted line for top panel and dashed-dotted line for bottom panel; 4th iteration: dotted line for bottom panel). Black arrows correspond to links existing in the adjacency matrix, i.e., direct links, and red arrows are purely hidden links absent from the adjacency matrix but present in the *G*_qr_ component of the reduced Google matrix *G*_R_. These networks have been drawn with Cytoscape [33].

It is worth to note that although *Leukemia* article in May 2017 English Wikipedia does not cite any of the other articles devoted to cancer types (as an illustration the first half of the *Leukemia* column in *G*_*rr*_ is filled with zero entries, see Fig. 5 top right panel), we are able to infer hidden links (in red in Fig. 6, top panel) from *Leukemia* to other cancers, here *Lung cancer* and *Non-Hodgkin lymphoma*.

In the reduced network of cancer, Fig. 6 (top panel), we connect to each cancer types the two preferentially linked countries, i.e., for each cancer type *cr*, the two countries *cn*_1_ and *cn*_2_ giving the two highest value (*G*_*rr*_ + *G*_qrnd_)_*cn,cr*_. We observe that cancers affecting digestive system point preferentially to Asian countries with the exception of Great Britain and Chile (*Liver cancer* points to Thailand and Saudi Arabia, *Stomach cancer* to Mongolia and Chile, *Colorectal cancer* to Philippines and Great Britain). This results are correlated to the fact that high mortality rates for *Liver cancer* are found in Asia (with the highest death rates for Eastern Asia [44]), and for *Stomach cancer* in Eastern Asia and South America [45, 46]. In the other hand *Colorectal cancer* epidemiology clearly states [47] that the highest incidence rates are found for Western countries such as Great Britain. The appearance of Philippines pointed by *Colorectal cancer* is an artifact due to the mention in the corresponding 2017 Wikipedia article of Corazon Aquino, former president of the Philippines who was diagnosed with this cancer type. Blood cancer types points preferentially to African countries with the exception of Cambodia pointed by *Hodgkin lymphoma*. At first sight this results can appear surprising since these blood cancers are found worldwide with incidence rates highest for Western countries and lowest for African countries [48]. In fact there is a Non-Hodgkin lymphoma, the Burkitt’s lymphoma [49], which mainly affects children in malaria endemic region, i.e., Equatorial and Sub-Equatorial Africa and Eastern Asia. Countries pointed by blood cancer types, i.e., Liberia, Zambia, Cameroon, Gabon and Cambodia, belong to these regions. Let us note that these cancers and countries are connected through hidden links. *Melanoma* points to Australia, which is, with New Zealand [50], the country having the highest rate of *Melanoma*, and points to Peru, where nine 2400 years old mummies have been found with apparent signs of *Melanoma* [50]. *Prostate cancer* points preferentially to Japan, due to its exceptional low incidence on Japanese population in Japan and abroad [51, 52], to Nigeria, since it is believe that black population is particularly at risk [53]. *Lung cancer* points to Germany, where in 1929 it was shown for the first time a correlation between smoking and *Lung cancer* [54, 55], and to Bhutan which adopted a complete smoking ban since 2005 [54]. Hidden link from *Breast cancer* to Republic of San Marino should be related to the fact that inhabitants of San Marino commemorate Saint Agatha, patroness of the Republic and of breast cancer patients [56]. Hidden link from *Breast cancer* to Cambodia is more difficult to interpret.

Let us now consider the reduced Google matrix associated to *N*_*r*_ = *N*_*cr*_ + *N*_*d*_ = 240 May 2017 English Wikipedia articles devoted to *N*_*cr*_ = 37 cancer types and to *N*_*d*_ = 203 cancer drugs. As above the reduced network of cancer can be constructed (Fig. 6, bottom panel). The construction process ends at the 4th iteration. The main structure of reduced network of cancers is the same as the previous with some exceptions. *Pancreatic cancer* is added to the digestive system cancers cluster and via hidden links, *Melanoma* points now to *Skin cancer* which points to *Breast cancer*. Consequently we observe a new cluster of thoracic region cancers comprising *Skin, Breast, Lung cancers* and *Melanoma*. Let us connect to each cancer types the two preferentially linked cancer drugs, i.e., for each cancer type *cr*, the two cancer drugs *d*_1_ and *d*_2_ giving the two highest value (*G*_*rr*_ + *G*_qrnd_)_*d,cr*_. Using DrugBank database [29], we easily check that indeed each drug is currently used to treat the cancer type to which it is connected. Also, closely connected cancer types share the same medication, e.g., *Skin cancer* and *Melanoma* are treated by *Vemurafenib* and *Dabrafenib* which are enzyme inhibitor of BRAF gene [57], *Leukemia* and *Non-Hodgkin lymphoma* are treated by the antibody *Rituximab* targeting B-lymphocyte antigen CD20 [58]. On the other hand non connected cancer types can in some cases share the same medication, the monoclonal antibody *Trastuzumab* typically used for *Breast cancer* is now also considered as a drug for *Stomach cancer* since these two cancer types overexpress the HER2 gene [59]. Let us note that hidden links connecting *Non-Hodgkin lymphoma* to *Cyclophosphamide* and *Rituximab* capture also a current medication reported in DrugBank database [29].

The reduced network of cancers shown in Fig. 6 depict in a relevant manner interactions between cancers, cancer-country and cancer-drug interactions through Wikipedia.

### World countries sensitivity to cancers

We consider the reduced Google matrix associated to the set of *N*_*r*_ = *N*_*cr*_ + *N*_*cn*_ = 232 Wikipedia articles constituted of *N*_*cr*_ = 37 articles devoted to cancer types and of *N*_*cn*_ = 195 articles devoted to countries. We compute the PageRank sensitivity *D*(*cr* → *cn, cn*), i.e., the infinitesimal rate of variation of PageRank probability *P* (*cn*) when the directed link *cr* → *cn*, (*G*_R_)_*cn,cr*_, is increased by an amount *δ*(*G*_R_)_*cn,cr*_, where *δ* is an infinitesimal.

Fig. 7 shows the world distribution of PageRank sensitivity *D*(*cr* → *cn, cn*) to *Lung cancer*. The most sensitive countries are, as discussed in the previous section, Bhutan and Germany mainly because these countries are directly cited in Wikipedia’s *Lung cancer* article. Besides articles devoted to these two countries the others are not directly linked from the *Lung cancer* article and the results obtained in Fig. 7 (top panel) is consistent with GLOBOCAN 2018 data [4]: apart Micronesia/Polynesia, the most affected countries, in term of incidence rates, are Eastern Europe, Eastern Asia, Western Europe, and, Southern Europe for males, and, Northern America, Northern Europe, Western Europe, and, Australia/New Zealand for females. The less affected are African countries for both sexes. Let us note that although incidence rates are very high for males in Micronesia/Polynesia according to [4], this fact is not captured by Wikipedia since Nauru, Kiribati, Tuvalu, Marshall Islands are the less PageRank sensitive countries. This is certainly due to the fact that articles devoted to these sovereign states are among the worst ranked articles devoted to countries in the May 2017 English Wikipedia ranking using PageRank algorithm. Their respective ranks are Nauru *K* = 7085, Kiribati *K* = 7659, Tuvalu *K* = 6201, Marshall Islands *K* = 4549 to compare e.g. with USA *K* = 1, France *K* = 4, Germany *K* = 5, etc (see PageRank indices of countries in [28]).

**Fig 7.**
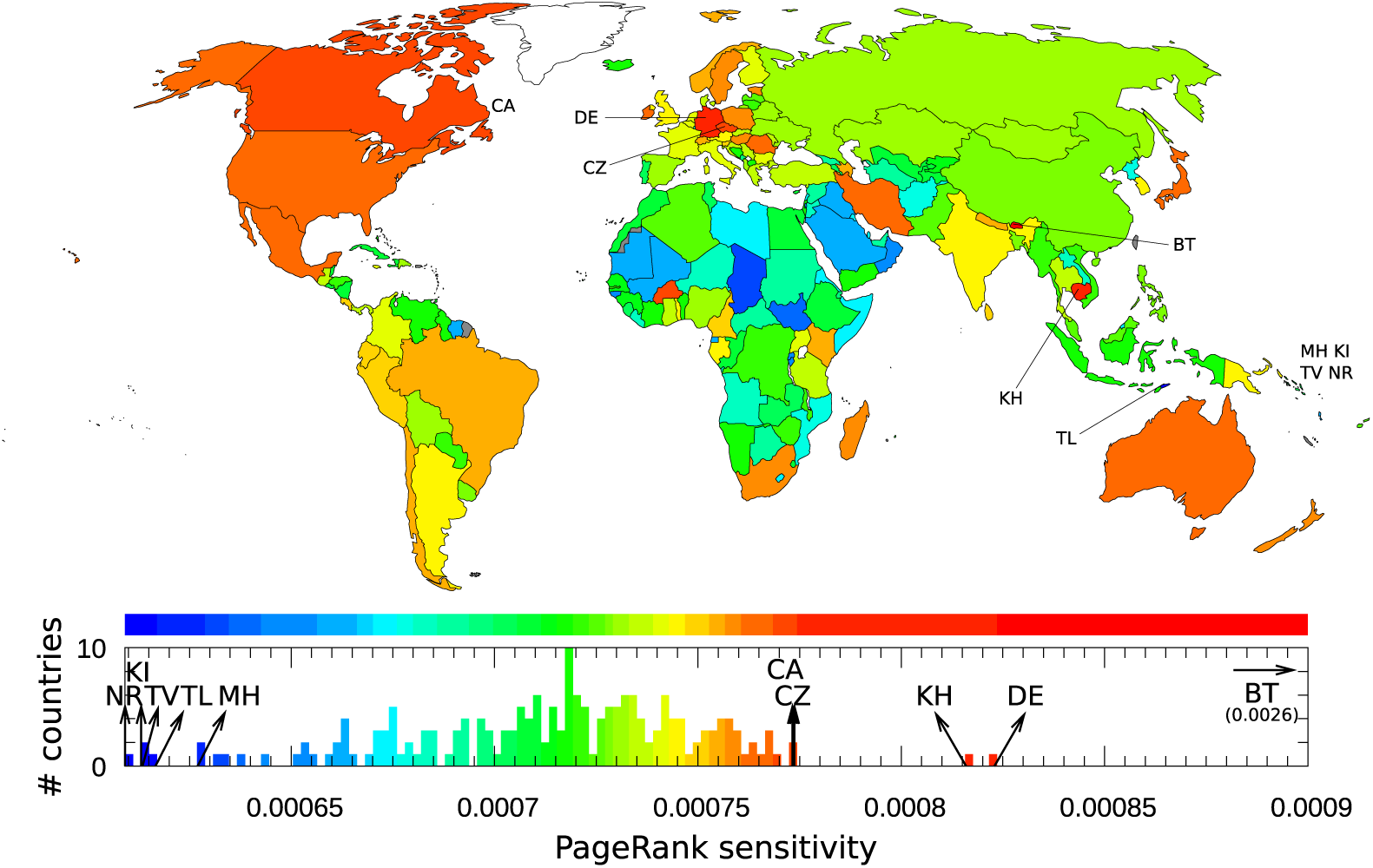
Sensitivity of countries to *Lung cancer*. A country *cn* is colored according to its diagonal PageRank sensitivity *D* (*cr* → *cn, cn*) to *Lung cancer*. Color categories are obtained using the Jenks natural breaks classification method [60].

As complementary information, sensitivities of countries to *Breast cancer* and to *Leukemia* are given in [28].

In order to investigate cancer – drug interactions it is also possible to represent sensitivity of countries to the variation of links from a cancer to a drug. As an illustration, Fig. 8 shows countries PageRank sensitivities to variation of *Lung cancer* → *Bevacizumab* link. We see that in this case the sensitivity of countries is significantly reduced comparing to the direct sensitivity influence of lung cancer on world countries shown in Fig. 7. Since the influence of this link variation is indirect for countries it is rather difficult to recover due to what indirect links the influence for specific countries is bigger or smaller. Among the most affected European countries we find Lichtenstein, Great Britain, Iceland, Portugal and Croatia while Germany and the Czech Republic are mostly unaffected. Another example of sensitivity of countries to cancer-drug link variation is given in [28].

**Fig 8.**
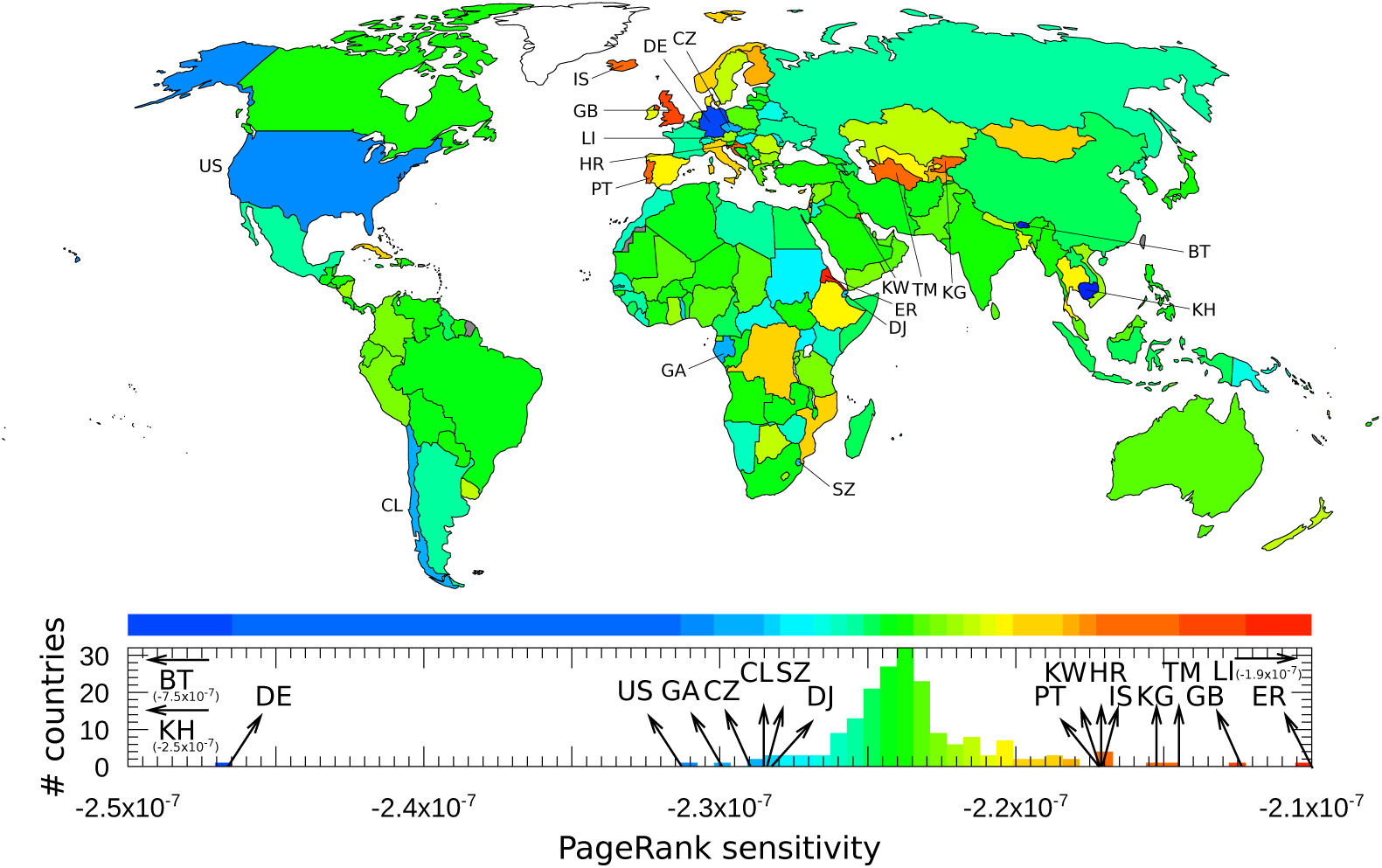
Sensitivity of countries to cancer → drug link variation. A country *cn* is colored according to its nondiagonal PageRank sensitivity *D* (*cr* → *d, cn*) to *cr* → *d* link variation. Variation of *Lung cancer* → *Bevacizumab* link is considered. Color categories are obtained using the Jenks natural breaks classification method [60].

### Interactions between cancers and drugs

Let us investigate interactions between cancers and drugs considering the subnetwork of *N*_*cr*_ = 37 cancers (see Tab. 1) and of the first 37 cancer drugs appearing in the PageRank ordered list Tab. 3. We do not consider *Talc* here since it is widely used in not only pharmaceutical industries.

We consider the sensitivity of cancer to drugs via the computation of *D* (*cr* → *d, cr*) presented in Fig. 9. Although the PageRank sensitivity is computed using the logarithmic derivative of the PageRank, globally the most sensitives cancers are the ones with the highest PageRank probability, i.e., the ones with lowest PageRank indices *K* (see Fig. 2 and Tab. 3): *Lung cancer* is mostly sensitive to *Irinotecan, Etoposide, Carboplatin, Breast cancer* to *Raloxifene, Trastuzumab, Docetaxel, Leukemia* to *Mercaptopurine, Imatinib, Rituximab*, etc. Following the National Cancer Institute [25] and/or DrugBank [29] databases, these associations cancer – drug are indeed approved.

**Fig 9.**
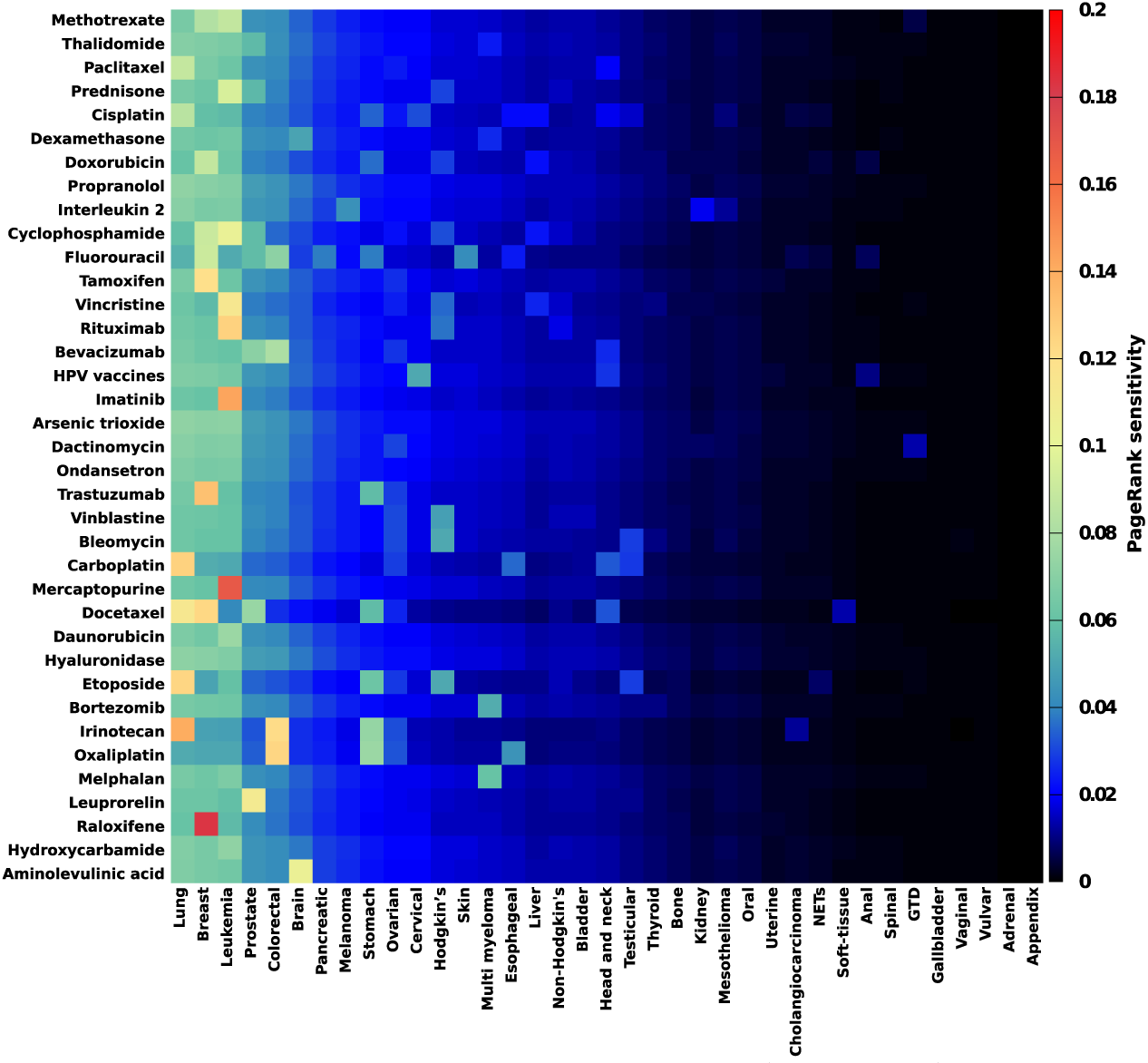
Sensitivity of cancers to drugs. The PageRank sensitivity *D* (*cr* → *d, cr*) of cancers to cancer drugs is represented. Here we consider the first 37 cancers (*cr*) listed in Tab. 3 and the first 37 drugs (*d*) listed in Table 2 (*Talc* has been removed as its article is too general).

Fig. 10 shows the complementary view of the sensitivity of drugs to cancers obtained from the computation of *D* (*d* → *cr, d*). Here the most sensitive drugs are *Dactinomycin* to *Gestational trophoblastic disease, HPV vaccines* to *Vulvar* and *Vaginal cancers, Fluorouracil* to *Anal cancer, Doxorubicin* to *Soft-tissue cancers*, etc. Again the National Cancer Institute [25] and DrugBank [29] databases report these possible drug – cancer associations.

**Fig 10.**
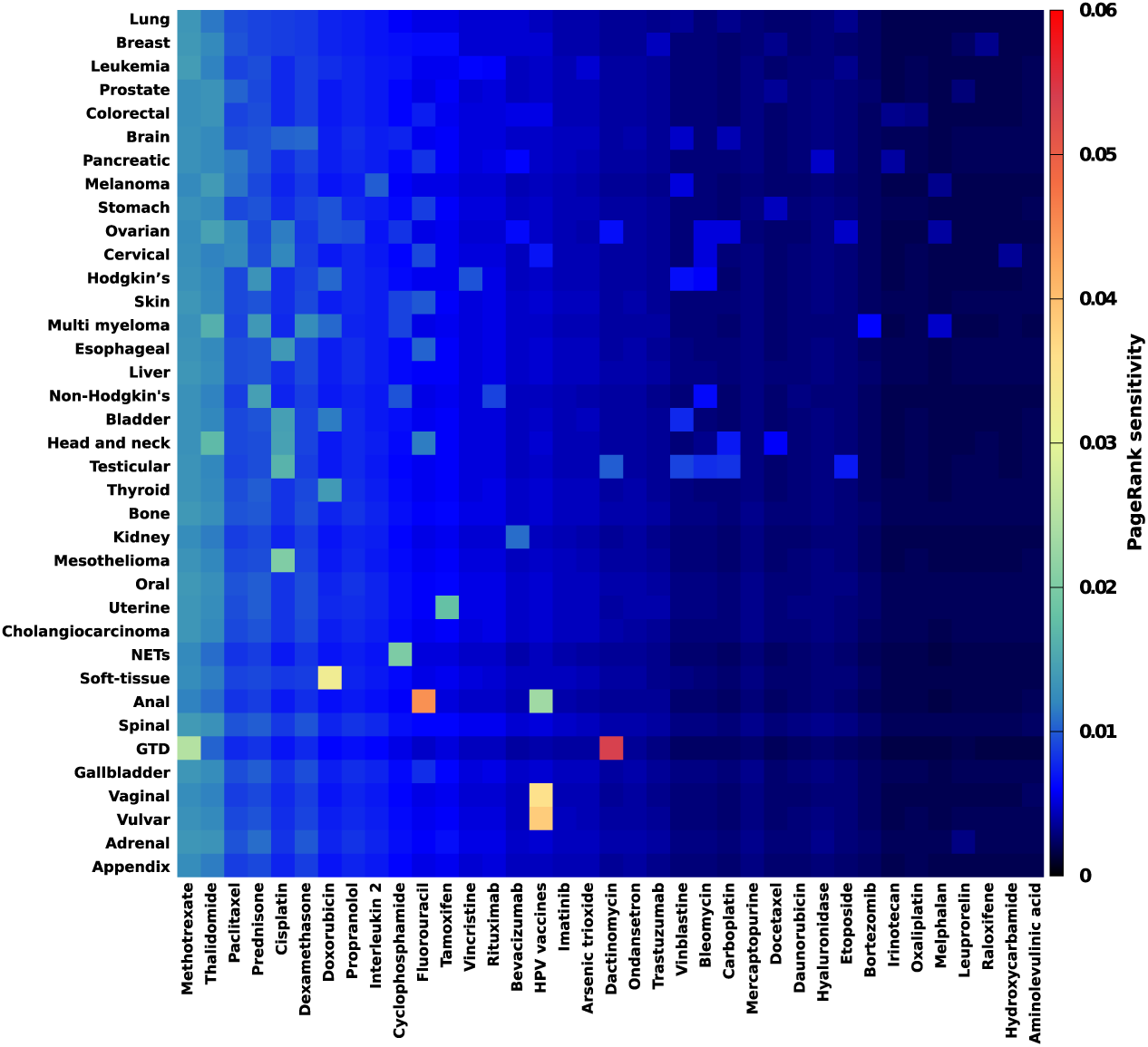
Sensitivity of drugs to cancers. The PageRank sensitivity *D* (*d* →*cr, d*) of cancer drugs to cancers is represented. Here we consider the first 37 cancers (*cr*) listed in Tab. 3 and the first 37 drugs (*d*) listed in Table 2 (*Talc* has been removed as its article is too general).

Let us consider directly the reduced Google matrix associated to the top 20 cancer types and top 20 cancer drugs according to May 2017 English Wikipedia PageRank list (Tab. 3). This reduced Google matrix *G*_R_ and its *G*_*rr*_, *G*_pr_ and *G*_qrnd_ components are shown in Fig. 5.

For each cancer *cr* of the 20 most influent cancer types in May 2017 English Wikipedia let us determine the three most connected drugs *d*, i.e., the three drugs with the highest value of (*G*_*rr*_ + *G*_qrnd_)_*d,cr*_. In Tab. 6 we show the May 2017 English Wikipedia prescription for each one of the top 20 cancer types. Most of the prescribed drugs are approved drugs for the considered cancer types according to National Cancer Institute [25] and DrugBank [29]. Some of the Wikipedia proposed drugs are in fact subject of passed, ongoing or planned clinical trials. Only Dexamethasone is in fact not specific to *Brain tumor* since it is a corticosteroid used to treat inflammation in many medical conditions. We observe that hidden links gives also accurate medication, see drugs associated to *Non-Hodgkin lymphoma* and *Bladder cancer* in Tab. 6.

**Table 6.** Drug prescription by Wikipedia for the top 20 most influential cancer types and comparison with prescriptions by National Cancer Institute and DrugBank. For each of the top 20 cancer types ranked in May 2017 English Wikipedia using PageRank algorithm (see Tab. 3), we give the three strongest cancer → drug links, i.e., for a given cancer type *cr* we select the three cancer drugs *d* with the highest values (*G*_*rr*_ + *G*_qr_)_*d,cr*_. Drug in red indicates a pure hidden cancer → drug link, i.e., the cancer type article in Wikipedia does not refer directly to the drug. For each cancer → drug link, the drug is followed by a ✓ mark if it is indeed prescribed for the cancer type according to National Cancer Institute [25] and/or DrugBank [29]; by a. ▴ mark if the drug appears only as a subject of passed, ongoing or planned clinical trials reported for the cancer type in DrugBank; and by a ✗ mark otherwise.

|  | Cancer | 1st drug | 2nd drug | 3rd drug |
| --- | --- | --- | --- | --- |
| 1 | Lung cancer | Erlotinib ✓ | Crizotinib ✓ | Cisplatin ✓ |
| 2 | Breast cancer | Tamoxifen ✓ | Trastuzumab ✓ | Methotrexate ✓ |
| 3 | Leukemia | Imatinib ✓ | Rituximab ✓ | Methotrexate ✓ |
| 4 | Prostate cancer | Enzalutamide ✓ | Cyclophosphamide ▲ | Prednisone ✓ |
| 5 | Colorectal cancer | Fluorouracil ✓ | Irinotecan ✓ | Bevacizumab ✓ |
| 6 | Brain tumor | Temozolomide ✓ | Dexamethasone $\times^a$ | Aminolevulinic acid ▲ |
| 7 | Pancreatic cancer | Fluorouracil ✓ | Gemcitabine ✓ | Protein-bound paclitaxel ✓ |
| 8 | Melanoma | Vemurafenib ✓ | Dabrafenib ✓ | Trametinib ✓ |
| 9 | Stomach cancer | Trastuzumab ✓ | Doxorubicin ✓ | Cisplatin ▲ |
| 10 | Ovarian cancer | Cisplatin ✓ | Tamoxifen ▲ | Bevacizumab ✓ |
| 11 | Cervical cancer | HPV vaccines ✓ | Cisplatin ✓ | Topotecan ✓ |
| 12 | Hodgkin's lymphoma | Prednisone ✓ | Cyclophosphamide ✓ | Vincristine ✓ |
| 13 | Skin cancer | Vemurafenib ✓ | Dabrafenib ✓ | Fluorouracil ✓ |
| 14 | Multiple myeloma | Dexamethasone ▲ | Elotuzumab ✓ | Bortezomib ✓ |
| 15 | Esophageal cancer | Cisplatin ✓ | Carboplatin ✓ | Fluorouracil ✓ |
| 16 | Liver cancer | Doxorubicin ▲ | Cisplatin ▲ | Sorafenib ✓ |
| 17 | Non-Hodgkin's lymphoma | Cyclophosphamide ✓ | Rituximab ✓ | Prednisone ✓ |
| 18 | Bladder cancer | Doxorubicin ✓ | Cisplatin ✓ | Methotrexate ✓ |
| 19 | Head and neck cancer | Cetuximab ✓ | Paclitaxel ✓ | Cisplatin ✓ |
| 20 | Testicular cancer | Etoposide ✓ | Cisplatin ✓ | Bleomycin ✓ |

Notes: <sup>a</sup> Dexamethasone may be used to decrease swelling around the tumor [61].

Conversely for each cancer drug *d* of the 20 most influent cancer drugs in 2007 English Wikipedia we determine the three most connected cancer types *cr*, i.e., the three cancer types with the highest value of (*G*_*rr*_ + *G*_qrnd_)_*cr,d*_. In Tab. 7 we show for which cancers a drug is prescribed according to May 2017 English Wikipedia. Again the results are globally in accordance with National Cancer Institute [25] and DrugBank [29] databases. We note that hidden links here correspond mainly to clinical trials, e.g., Imatinib is an approved drug for treatment of certain forms of *Leukemia*, but experiments were or will be done for *Breast cancer* and *Prostate cancer*.

**Table 7.** According to Wikipedia for which cancer type is prescribed the top 20 most influential cancer drugs and comparison with prescriptions by National Cancer Institute and DrugBank. For each of the top 20 cancer drugs ranked in May 2017 English Wikipedia using PageRank algorithm (see Tab. 3), we give the three strongest drug → cancer links, i.e., for a given drug *d* we select the three cancer types *cr* with the highest values (*G*_*rr*_ + *G*_qr_)_*cr,d*_. Cancer type in red indicates a pure hidden drug → cancer link, i.e., the drug article in Wikipedia does not refer directly to the cancer type. For each drug →cancer link, the cancer type is followed by a ✓ mark if the drug is indeed prescribed for the cancer type according to National Cancer Institute [25] and/or DrugBank [29]; by a. ▴ mark if the drug appears only as a subject of passed, ongoing or planned clinical trials reported for the cancer type in DrugBank; and by a ✗ mark otherwise.

|  | Drug | 1st cancer type | 2nd cancer type | 3rd cancer type |
| --- | --- | --- | --- | --- |
| 1 | Talc | Ovarian cancer × | Lung cancer × | Breast cancer × |
| 2 | Methotrexate | Leukemia ✓ | Breast cancer ✓ | Lung cancer ✓ |
| 3 | Thalidomide | Multiple myeloma ✓ | Breast cancer ▲ | Prostate cancer ▲ |
| 4 | Paclitaxel | Breast cancer ✓ | Lung cancer ✓ | Ovarian cancer ✓ |
| 5 | Prednisone | Multiple myeloma ▲ | Non-Hodgkin lymphoma ✓ | Hodgkin's lymphoma ✓ |
| 6 | Cisplatin | Lung cancer ✓ | Testicular cancer ✓ | Breast cancer ✓ |
| 7 | Dexamethasone | Multiple myeloma ▲ | Brain tumor $\times^a$ | Leukemia ✓ |
| 8 | Doxorubicin | Leukemia ✓ | Hodgkin's lymphoma ✓ | Breast cancer ✓ |
| 9 | Propranolol | Ovarian cancer ▲ | Brain tumor × | Colorectal cancer ▲ |
| 10 | Interleukin 2 | Melanoma ✓ | Leukemia ▲ | Hodgkin's lymphoma ▲ |
| 11 | Cyclophosphamide | Leukemia ✓ | Multiple myeloma ✓ | Breast cancer ✓ |
| 12 | Fluorouracil | Colorectal cancer ✓ | Breast cancer ✓ | Stomach cancer ✓ |
| 13 | Tamoxifen | Breast cancer ✓ | Uterine cancer ▲ | Prostate cancer ▲ |
| 14 | Vincristine | Leukemia ✓ | Hodgkin's lymphoma ✓ | Lung cancer ✓ |
| 15 | Rituximab | Leukemia ✓ | Non-Hodgkin lymphoma ✓ | Multiple myeloma ▲ |
| 16 | Bevacizumab | Breast cancer ✓ | Colorectal cancer ✓ | Lung cancer ✓ |
| 17 | HPV vaccines | Cervical cancer ✓ | Breast cancer × | Colorectal cancer × |
| 18 | Imatinib | Leukemia ✓ | Breast cancer ▲ | Prostate cancer ▲ |
| 19 | Arsenic trioxide | Leukemia ✓ | Brain tumor ▲ | Breast cancer ▲ |
| 20 | Dactinomycin | GTD<sup>b</sup> ✓ | Testicular cancer ✓ | Ovarian cancer ✓ |

Notes: <sup>a</sup> Dexamethasone may be used to decrease swelling around the tumor [61]. <sup>b</sup> Gestational trophoblastic disease.

It would be interesting to thoroughly study the most connected drugs and cancer types from the hidden contributions only, i.e., from (*G*_qrnd_)_*cr,d* or *d,cr*_ matrix elements only, in order to test the possible predictive power of the reduced Google matrix analysis of Wikipedia networks. This study is beyond the scope of the present work and will be considered in a subsequent one.

## Conclusion

Using PageRank and CheiRank algorithms, we investigate global influences of 37 cancer types and 203 cancer drugs through the prism of Human knowledge encoded in the English edition of Wikipedia considered as a complex network. From the ranking of Wikipedia articles using PageRank algorithm we extract the ranking of the most influent cancers according to Wikipedia. This ranking is in good agreement with rankings, by either mortality rates or yearly new cases, extracted from WHO GLOBOCAN 2018 [2] and Global Burden of Diseases study 2017 [5] databases.

The recently developed algorithm of the reduced Google matrix allows to construct a reduced network of cancers taking into account all the information aggregated in Wikipedia. This network exhibits direct and hidden links between the most influent cancers which form clusters of similar or related cancer types. The reduced Google matrix gives also countries or cancer drugs which are preferentially linked to the most influent cancers. Inferred relations between cancer types and countries obtained from Wikipedia network analysis are in accordance with global epidemiology literature. The PageRank sensitivity of countries to cancer types gives also a complementary tool corroborating epidemiological analysis. As far as we know, it is the first study highlighting correspondence between Wikipedia network analysis and disease burden or epidemiological studies. Inferred interactions between cancers and cancer drugs allows to determine drug prescriptions by Wikipedia for a specific cancer. These Wikipedia prescriptions appear to be compatible with approved medications reported in National Cancer Institute [25] and DrugBank [29] databases.

The reduced Google matrix algorithm allows to determine a clear and compact description of global influences and interactions of cancer types and cancer drugs integrating well documented medical aspects but also historical, and societal aspects, all encoded in the huge amount of knowledge aggregated in Wikipedia since 2001.

## Authors contributions

All the authors were involved in the preparation of the manuscript. All the authors have read and approved the final manuscript.

## Acknowledgments

We thank Jean-Paul Feugeas and Tatiana Serebriyskaya for useful remarks and discussions. This work was supported by the French “Investissements d’Avenir” program, project ISITE-BFC (contract ANR-15-IDEX-0003), by the Bourgogne Franche-Comté Region 2017-2020 APEX project (conventions 2017Y-06426, 2017Y-06413, 2017Y-07534; see http://perso.utinam.cnrs.fr/~lages/apex/ and in part by the Programme Investissements d’Avenir ANR-11-IDEX-0002-02, reference ANR-10-LABX-0037-NEXT (project THETRACOM).

